# A hybrid stochastic-deterministic mechanochemical model of cell polarization

**DOI:** 10.1101/785709

**Authors:** Calina Copos, Alex Mogilner

**Affiliations:** Courant Institute, New York University, New York, NY, United States of America; Courant Institute and Department of Biology, New York University, New York, NY, United States of America

## Abstract

Polarization is a crucial component in cell differentiation, development, and motility and its details are not yet well understood. At the onset of cell locomotion, cells break symmetry to form a well-defined cell front and rear. This polarity establishment varies across cell types: in *Dictyostelium discoideum* cells, it is mediated by biochemical signaling pathways and can function in the absence of a cytoskeleton, while in keratocytes it is tightly connected to cytoskeletal dynamics and mechanics. Theoretical models that have been developed to understand the onset of polarization have explored either signaling or mechanical pathways, yet few have explored mechanochemical mechanisms. However, many motile cells rely on both signaling modules and actin cytoskeleton to break symmetry and achieve a stable polarized state. We propose a general mechanochemical polarization model based on the coupling between a stochastic model for the segregation of signaling molecules and a simplified mechanical model for actin cytoskeleton network competition. We find that local linear coupling between minimally nonlinear signaling and cytoskeletal systems, separately not supporting stable polarization, yields a robustly polarized cell state.

## Introduction

The ability to spontaneously break symmetry is fundamental to most eukaryotic cells and plays an important role in embryogenesis, cell differentiation, cell division, and migration. Intrinsically motile cells can spontaneously switch to a migratory polarized phenotype [47]. Understanding complex molecular circuits employed by a cell to establish polarization has been studied both theoretically [2, 19, 31, 32, 36, 42] and experimentally [6, 45, 49, 70].

Polarity establishment arises primarily through the localization of particular proteins and lipids in the cell to specific regions of the plasma membrane, and often precedes motility. Experiments have identified a few conserved sets of proteins involved in polarization including the PAR system [37, 44], the Wnt system [28], the Scribble complex [3, 58], and the Rho system [6, 55]. Here, we focus on the Rho molecular circuit whose dynamics can lead to cell polarization at the onset of cell motility. The Rho family of GTPases is a family of small proteins that act as molecular switches [22, 55]. Three important members of the family have been studied in detail: Cdc42, Rac1, and RhoA [55]. These proteins cycle between an inactive (GDP) cytosolic and an active (GTP) membrane-bound form that signals to the actin cytoskeleton and other downstream targets. Mutual antagonistic interactions between Rac1 and RhoA were identified, as well as spatial and/or temporal exclusions that produce a tendency for them to segregate to the front versus rear of a polarized cell [6, 7, 65, 72]. From previous theoretical work, it is well known that mutually inhibitory circuits, like those in Rac1/RhoA, could yield a robustly polarized system [14, 29].

Cell polarization is also associated with the rearrangement of the actin cytoskeleton during polarization, branched actin filaments form at the cell front while actomyosin contractile bundles segregate to the cell rear [60, 70, 73]. Just as diffusible chemical activators and inhibitors trigger biochemical instabilities, mechanical instabilities can arise due to stochastic fluctuations in actin filament densities or mechanical feedback between motor proteins and cytoskeletal elements [66]. In mechanically-driven polarity systems, cells polarize due to forces and actin flow generated by these forces [4, 18, 32, 43, 73]. Two classic cases involving cytoskeleton-driven polarization are the formation of actin comet tails by intracellular pathogens [12, 21] and the directional locomotion of keratocytes [4, 32, 73]. In both cases, the mechanical properties of the actin cytoskeletal network appear sufficient for polarization, which can be triggered by stochastic or induced asymmetries in the mechanical network.

Mathematical models have been used to explain spontaneous pattern formation in cells since the 1950s [38, 63]. Initial approaches were based on Turing patterns and focused on biochemical signaling pathways for polarity. In Turing-like models, chemical patterns emerge from stochastic fluctuations combined with interactions between chemical species that diffuse at different rates; these models often require elaborate nonlinearities for stable polarized distributions of chemicals [26, 29].

Recently, many models for cell polarization were proposed based on reaction-diffusion (not necessarily Turing-like) equations [29]. In one of the most popular models, the ‘wave-pinning’ model (Fig. 1 a), a minimal bistable reaction-diffusion system gives rise to polarization of the active/inactive chemicals [42]. In the model, the active form of the chemical diffuses slowly on the membrane and autocatalytically activates the inactive form of the chemical that diffuses fast in the cytoplasm. Conservation of the total chemical and the fast diffusion of its inactive form act as global inhibitors and pin or ‘arrest’ the active chemical in space into a stable polarized distribution. The model reproduces a number of observed features shared by many eukaryotic cells: (a) spontaneous (self-)polarization, (b) maintenance of polarized state after a stimulus is removed, (c) sensitivity to new incoming signals and ability to re-polarize in a new direction.

**Figure 1:**
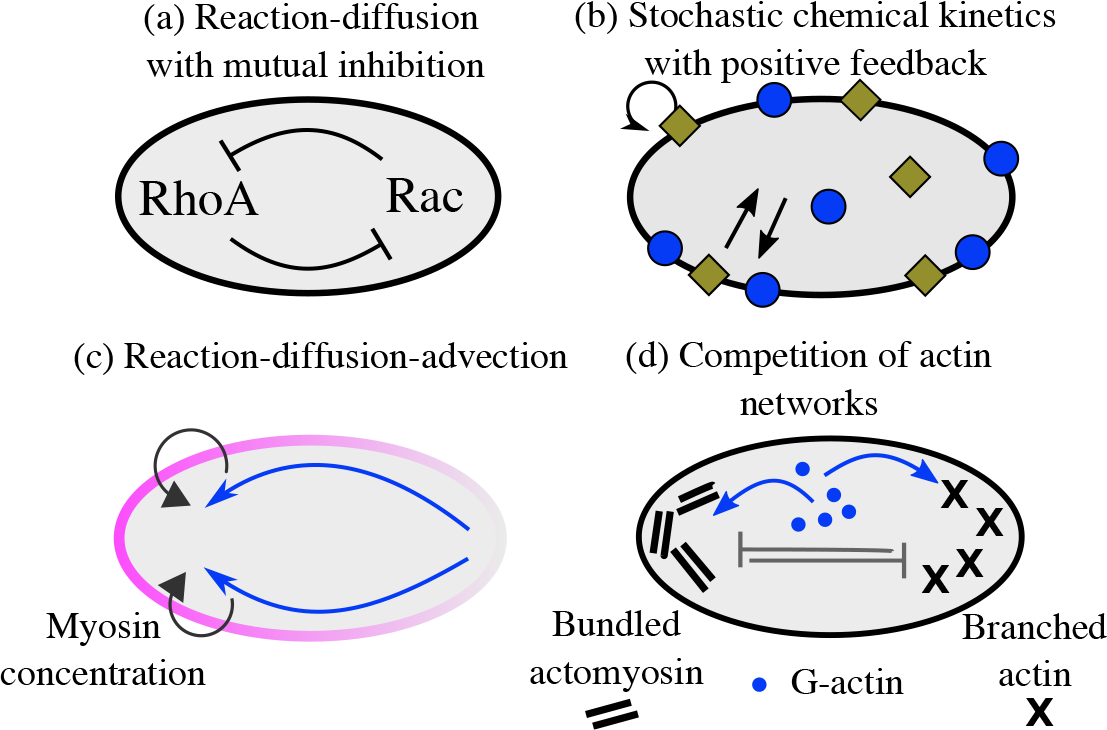
Simple models of cell polarization. Biochemical (a–b) and mechanical (c–d) approaches for polarity establishment. (a) Reaction-diffusion model of Rho GTPases signaling pathways are based on the mutual antagonism between signaling molecules. (b) Stochastic model of the cycling of active-inactive states of signaling molecules with autocatalytic feedback. (c) Polarization can also emerge due to positive feedback between myosin concentration gradients, which generates actomyosin flow up the gradient, and myosin transport by the flow. (d) In an advection-free model, polarization emerges due to the competition between branched protrusive actin network and contractile bundles of actomyosin network caused by competition for a conserved number of molecular resources.

Besides the reaction-diffusion models, stochastic polarization models have also been proposed (Fig. 1 b) [2, 46, 71]. For example, Altschuler et al. [2] found that clusters of active membrane-associated molecules can form and persist in time if there is a positive feedback loop in which active molecules recruit additional copies of itself from a cytoplasmic pool, provided the system is operating within a stochastic regime and the molecule number is limited.

A much smaller body of literature exists for the actin-driven models for cell polarization [4, 18, 32, 64]. Most of these models focus on the fast-moving fish epithelial keratocytes, which do not require the stereotypical signaling cascades to polarize [53]. A combined experimental and theoretical effort showed that the mechanical feedback between actin network flow, myosin, and adhesion is sufficient to amplify stochastic fluctuations in actin flow and trigger polarization (Fig. 1 c); aggregation of myosin at the cell rear generates rearward actin flow and forward cell movement, which both amplify the myosin concentration at the rear [4]. An actin flow-free mechanical mechanism for cell polarization was proposed by Lomakin et al. (Fig. 1 d) [32]. The authors demonstrated that the competition between branched and bundled actin networks around the cell periphery leads to the segregation of the actin cytoskeleton into branched filaments at the cell front and actomyosin bundles at the rear. A key assumption in this model is that protrusion of the boundary favors the branched network, while boundary retraction favors the contractile bundles.

Deterministic models for biochemical polarization mechanisms have limitations. For example, in the wave-pinning model, the stability of the polar distribution of the active chemical requires highly nonlinear reaction terms [42]. Furthermore, when the initial condition in the wave-pinning model consists of multiple localized patches of active chemicals, and when the diffusion constant is very small, these initial patches persist in time corresponding to formation of not one but multiple zones of activity on the plasma membrane (Fig. 2 a). Stochastic models for polarity molecules require very simple kinetics, but also require additional assumptions to constraint the number and location of emergent clusters into a single one (Fig. 2 d, *N* = 100) [2].

**Figure 2:**
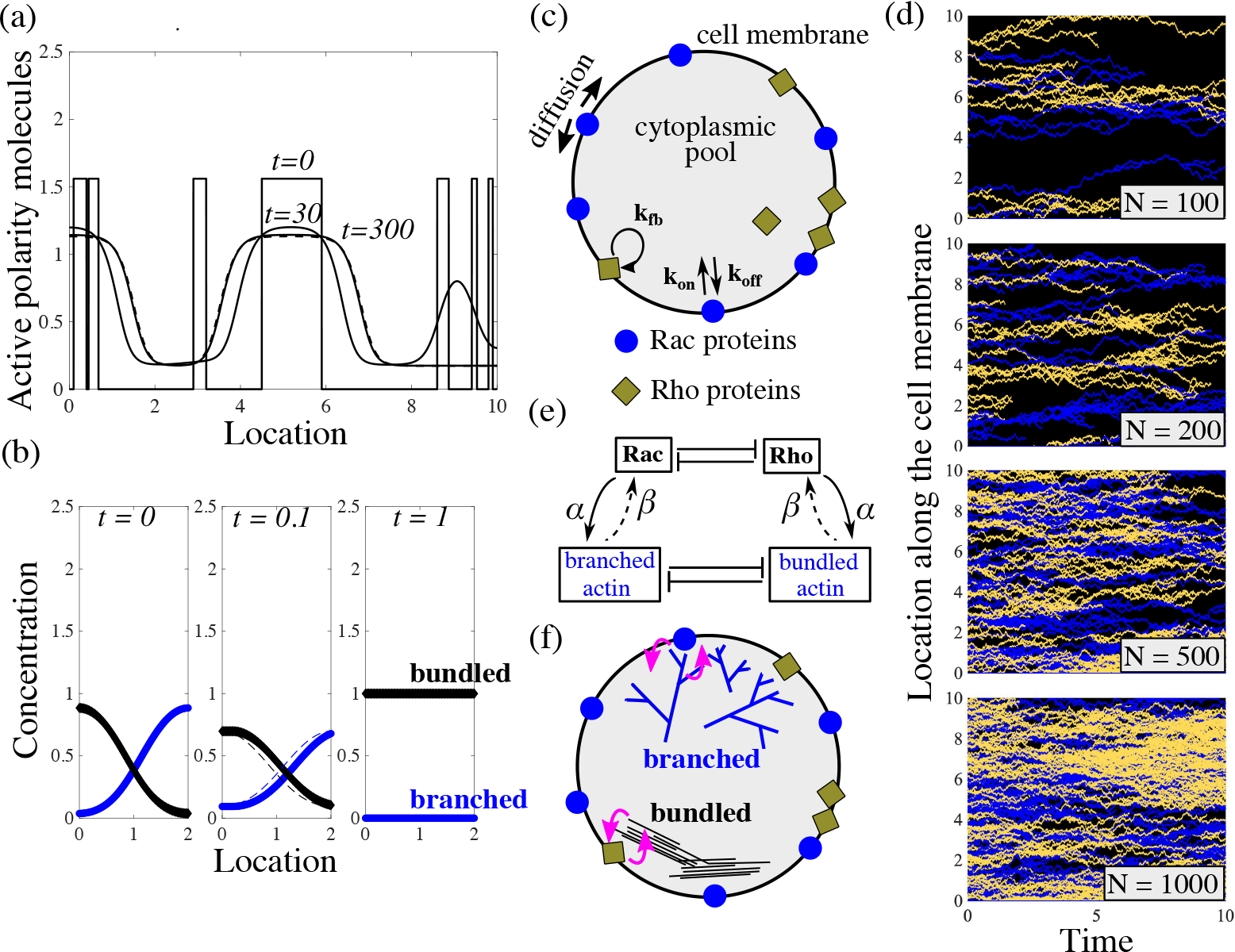
Neither model produces a stable polarized cell state. (a) Given ‘patchy’ initial conditions and low diffusion on the membrane, the wave-pinning model [42] cannot yield a polar distribution of the membrane-bound active polarity molecules. (b) In the mechanical cytoskeleton model for polarization of keratocytes without movement [32], perturbations to the polar distributions of the branched (*A*) and bundled (*B*) actin networks lead to annihilation of one of the networks. (c) Schematic of the stochastic polarization model modified from Altschuler et al. [2] describes the cycling of active-inactive states of two species of signaling molecules: Rac1 and RhoA. (d) Results of model simulations from (c) with varying total number of a species *N*. With a small number of particles, long-lasting patches of active form of Rac and Rho appear, however there is no control on the number of emergent patches or their merging into single protrusive and contractile fronts. Decreasing the number of signaling molecules leads to increasing levels of spatial segregation between the two signaling species. (e) In our coupled model, we link the non-polarizing mechanical model for the cytoskeleton to the non-polarizing simple kinetics model for Rac/Rho dynamics. (f) Schematic of the coupled model: Rac proteins and the branched actin network engage in mutual local feedback and similarly, so do Rho proteins and the bundled actin network.

Although cell polarity can emerge from systems that are either chemical or mechanical, in many cases, cell polarity depends on the interplay between the two [5, 11, 26, 51]. One such example of mechanochemical polarization is the establishment of the anterior-posterior axis in *Caenorhabditis elegans* embryos, which depends on both actomyosin flow and the biochemistry of PAR polarity proteins [11, 18, 44]. In this system, asymmetrical cell contractility and cortical actin flow are essential for polarity establishment [11, 44]. By coupling an advective transport of the flowing cell cortex to a reaction-diffusion system for PAR protein segregation, Goehring et al. [18] showed that advection could serve as a mechanical trigger and would be sufficient to form stable asymmetric PAR distribution. Similar experimental observations of the feedback between mechanics and biochemical signaling in polarity of other organisms continue to appear [20].

Here, we set out to uncover a minimal coupling between the simplest biochemical signaling and cytoskeletal circuits that support robust cell polarization. The competition between two actin networks (branched and protruding vs. bundled and contracting) is one such minimal cytoskeletal mechanism not requiring advective actin flow [32]. However, to generate stable polarization, this mechanism requires coupling to physical cell movement. Most cells move slowly and break symmetry before initiating locomotion [51]. Here, we show that the simplest feedback between the two-network competition and the simplest stochastic model of biochemical polarization with minimal nonlinearities leads to robust cell polarization.

## Model Formulation

### Minimal biochemical signaling model

The biochemical part of our model is a caricature of well-studied Rho GTPases kinetics [40]. Each Rho GTPase molecule cycles between two states: an active GTP-bound form, bound to the plasma membrane, and an inactive GDP-bound form, diffusing in the cell cytosol [6, 22]. The conformational changes between these two states are facilitated by a class of regulatory proteins like GEFs, GAPs, or GDIs [6, 22]. Rac and RhoA pathways crosstalk with one another, and previous research has revealed evidence of mutual inhibition of Rac and Rho signaling [6, 7, 65, 72]. Furthermore, this mutual antagonism is believed to promote spatial and/or temporal exclusions that produce a tendency for Rac and RhoA segregation [74].

At the onset of polarity, nascent polar sites appear along the plasma membrane and their presence is believed to be sustained by local positive feedback that depends on the assembly of branched actin filaments [27, 33]. These initial polar sites are thought to be sustained through local positive feedback loops that depend both the presence of polarity molecules [45], but also actin assembly [27, 57, 69]. Specifically, in their active state, Rho GTPases can bind to downstream effector proteins that control the actin cytoskeleton rearrangements [23, 52]. Rac polarity sites mediate the formation of a branching actin filament network at the leading edge [68]. Interaction between Rac and branched actin network appears to be mutual – Rac activates nucleation-promoting factors such as WAVE and WASP, which activate Arp2/3 branching complexes [24, 34], while recruitment of additional Rac in nascent polarity zones in highly protrusive regions is also reported [10, 45, 67]. RhoA is believed to stimulate the formation of a bundled actomyosin network through recruitment of myosin-II molecular motors at the opposing end [22, 50]. While less is known about the interaction from actomyosin bundles to RhoA, recent work seems to indicate that localization of RhoA activators is actin dependent [56].

Our signaling kinetics model is formulated on a one-dimensional (1D) computational domain representing polarity molecules on the plasma membrane and a thin volume of cytoplasm adjacent to the membrane on the circular edge of a disc-like cell spread on a flat surface. Position of the molecules is represented by the arc length *s* on a circle. We track coordinates of activated Rac and Rho molecules on the membrane along the cell edge, 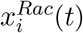 and 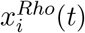, where *i* is the index of the specific molecule. Cytoplasmic concentrations of *i i* Rac and Rho are assumed to be homogeneous due to the fast diffusion in the cytoplasm. For the dynamics of these proteins, we extend the theory of Altschuler et al. [2] to include two species of signaling molecules. The redistribution of signaling molecules is determined by the rates of four mechanisms (Fig. 2c): (1) positive-feedback-induced activation and recruitment of cytoplasmic molecules to the locations of membrane-bound active signaling molecules with rate *k*_fb_; (2) spontaneous activation and association of cytoplasmic molecules to random locations on the plasma membrane with rate *k*_on_; (3) lateral diffusion (with coefficient *D*) of active molecules along the membrane; and (4) deactivation and disassociation of signaling molecules from the membrane with rate *k*_off_. For simplicity, we use the same kinetic rates for both Rac and Rho. Based on these dynamics, the number *n*^Rac^(*t*) of Rac molecules on the membrane at time *t* evolves by the continuous time

Markov process provided by the following master equation:

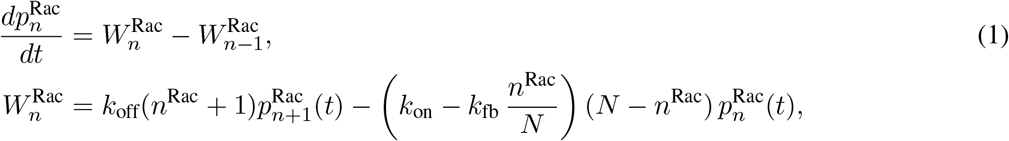

where 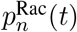 is the probability of having *n*^Rac^ Rac molecules on the membrane. Similar master equation with the same baseline kinetic rates is used for Rho molecules. The model parameters *k*_fb_, *k*_on_, *D* and *k*_off_, and total number of signaling molecules *N*, are readily biologically interpretable, and may be estimated from experimental data (see Table S1). In the diffusion process, a steric repulsion is enforced between Rac and Rho polarity molecules so that the two chemicals cannot spatially overlap at any moment in time. The rates of transition [2, 10, 16, 41, 42, 75], relative rates of diffusion [39], and concentrations of active and inactive states [2, 10] are known. In the Supplemental Material, we explain in detail how the diffusive random walk of the molecules on the membrane is simulated, as well as the numerical implementation of the model.

### Minimal actin network model

Two different actin structures are characteristic of polarized cell migration: a branched, protrusive actin network at the cell front, and contractile network made up of actomyosin bundles at the cell rear [32]. These two actin networks compete mechanically [32]: protrusion of the branched network leaves the bundled network behind the cell edge, while the contractile bundles collapse the branched filaments into bundles. The networks also compete for the same pool of actin monomers and other molecular resources in the cytosol [54]. Thus, our model has the features of a system with two competing species [13]. The spatiotemporal distributions of the two species of actin networks along the plasma membrane arc length are *A*(*s*, *t*) for the protrusive network and *B*(*s*, *t*) for the contractile actin-myosin meshwork. We assume that the branched network is protrusive and devoid of myosin-II motors, while the bundled network contains contractile actomyosin bundles that generate contractile forces and retract the cells posterior. Their dynamics are given by the following non-dimensionalized coupled system of equations adapted from Lomakin et al. [32]:

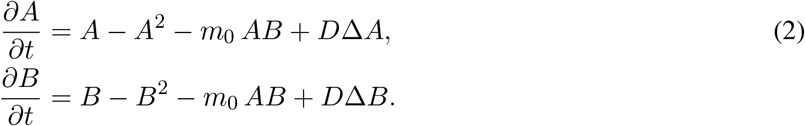

Full model assumptions and details are explained in the Supplemental Material. The rate of the network growth is proportional to their density, but it is limited at high densities by the finite amounts of molecular resources (e.g. depletion of monomers or branching complexes or myosin-II motors). The model assumes that the competition terms, proportional to the product *AB*, stem from either mechanical competition or competition for limited molecular resources. Parameter *m*_0_ is the magnitude of the competition between the networks. Lastly, the diffusive terms describe the action of myosin-II motors that slide and shuffle bundled filaments in the contractile actomyosin network, as well as the random lateral displacements of the growing ends of the barbed filaments along the cell membrane [32].

### Mechanochemical coupling

To couple the cytoskeleton meshwork dynamics to the cycling of signaling molecules, we assume, based on the experimental evidence described above, the simplest possible local feedback loops between (1) Rac and branched actin network, and (2) Rho and bundled actin network (Fig. 2f). Specifically, we posit that the chemical rates in the signaling module are no longer constant but rather are linear functions of the local concentration of each respective actin network which evolves in space and time. On the reverse, the growth rate of each actin network is linearly proportional to the respective local densities of active polarity proteins. We use the following mathematical expressions for the modified rates of Rac and Rho kinetics (Fig. 2e):

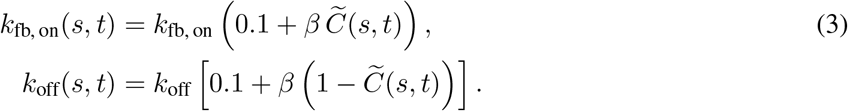

Here, the expressions for on and off rates are either for Rac or Rho chemical kinetics. For Rac/Rho kinetics, 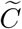 denotes branched/bundled actin network density, *A/B*, respectively, normalized by its largest value at any given time point. Thus, the induced on rate for Rac increases with the local branched actin density, while the Rac off rate decreases with the local branched actin density. Similar dependencies hold for the association and disassociation rates of Rho kinetics on the local bundled actin density. Coupling constant *β* is the measure of this feedback strength. The growth term in the actin networks equations is altered as follows:

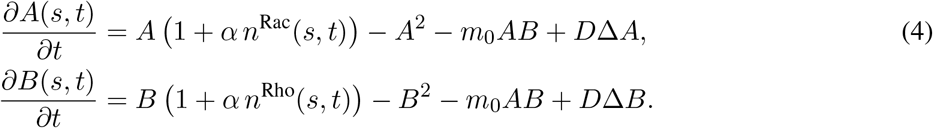

The coupling constant *α* is the measure of the feedback from Rac/Rho to the branched/bundled actin, respectively. In these expressions, *n*^Rac^(*s*, *t*) and *n*^Rho^(*s, t*) are densities of the Rac and Rho, respectively. Numerically, these densities are computed from the discrete locations of the respective molecules by computing at each time step a superposition of Gaussian peaks with variance *ε* and centers at their molecular locations. Note that in the presence of the feedback from actin networks to the signaling molecules, their on/off rates depend on positions on the cell edge due to varying actin densities. In this case, simulations of the chemical kinetics becomes more complex than solving Eq. 1 and numerical implementation of this modified stochastic model is described in the Supplemental Material. The codes used are accessible online. The numerical value for the coupling constants *α* and *β* ranges between 0 (uncoupled) and 1 (fully coupled).

## Results

### Neither the mechanical nor the biochemical model leads to stable polarization

A deterministic formulation of the polarity signaling model in Fig. 1c with only one chemical species, results in a reaction-diffusion equation with stably non-polar distributions [2]. On the other hand, a stochastic formulation of the same kinetics leads to the emergence of clusters of Rac and Rho that persist in time (Fig. 2b). However, we find that neither Rac, nor Rho clusters localize into a single cluster; Rac and Rho do not concentrate to opposite sides of the polarized cell, as observed at the onset of motility. As the total number of signaling molecules is increased, the number of clusters of Rho GTPases activity also increases resulting in a ‘patchy’ distribution. We varied the biochemical kinetic rates and found that these results hold over a wide range of model parameters.

The actin competition model is based on the classical Lotka-Volterra model of the population dynamics of two species competing for a common resource [13]. If the competition parameter *m*_0_ is small, two networks can coexist in space, and both of their densities are spatially constant. However, if the competition parameter *m*_0_ is large, the situation becomes ‘winner takes all’, and one network dominates, while the other goes extinct. We examined if polarized actin distributions are possible: one of the networks occupies one region in space, while the other occupies the remaining space (Fig. 2b). Such polarized state exists when the initial distributions are exactly symmetric. Any small perturbation of the initial polarized distribution results in one network (the one with slightly greater spatial support) taking over and displacing the other network (Fig. 2b). Rigorous mathematical analysis showed that this nontrivial spatial segregation of competing species in Lotka-Volterra models is not stable [8, 30, 61].

These simulations led us to wonder if coupling between the chemical and mechanical modules could stabilize cell polarization. The idea is that the slow destabilitization of the polar state of actin networks could bias the signaling molecules into segregating to two opposite parts of the cell; this bias is caused by the up-regulation of Rac on the membrane by the branched network and simultaneously, the up-regulation of Rho by the bundled network. Then, patches of Rac and Rho, whose positions are arbitrary without feedback, could drift to opposing cell regions. In turn, feedback from Rho GTPases to the respective actin networks ensures that one network does not invade another’s territory, since each network dominates in a specific region due to the presence of ‘its supporting’ chemical.

### The coupled mechanochemical model produces symmetry breaking depending on coupling strength

To assess whether this model with positive feedback between protrusive actin and Rac, and contractile actin and Rho, can account for symmetry breaking, we run simulations with two choices for the coupling constants (Fig. 3). The numerical simulations assume random initial distributions and equal total conserved amounts of Rac and Rho, and of two types of actin network (Fig. 3 a,e). For low coupling constants, the polarity proteins segregated into many non-overlapping Rac and Rho clusters (Fig. 3d). The mechanical model was unable to achieve a polar distribution: either the actomyosin bundles or the branched actin meshwork went extinct (Fig. 3c). In this particular instance of the simulation, the bundled actomyosin meshwork died out while the protrusive network survives and takes over the entire plasma membrane (Fig. 3c). Tens of instances for this set of parameters were considered and a probability of polarization establishment of 30% was computed as a fraction of number of simulations with stable polar distributions of both actin and polarity chemicals over the total number of runs.

**Figure 3:**
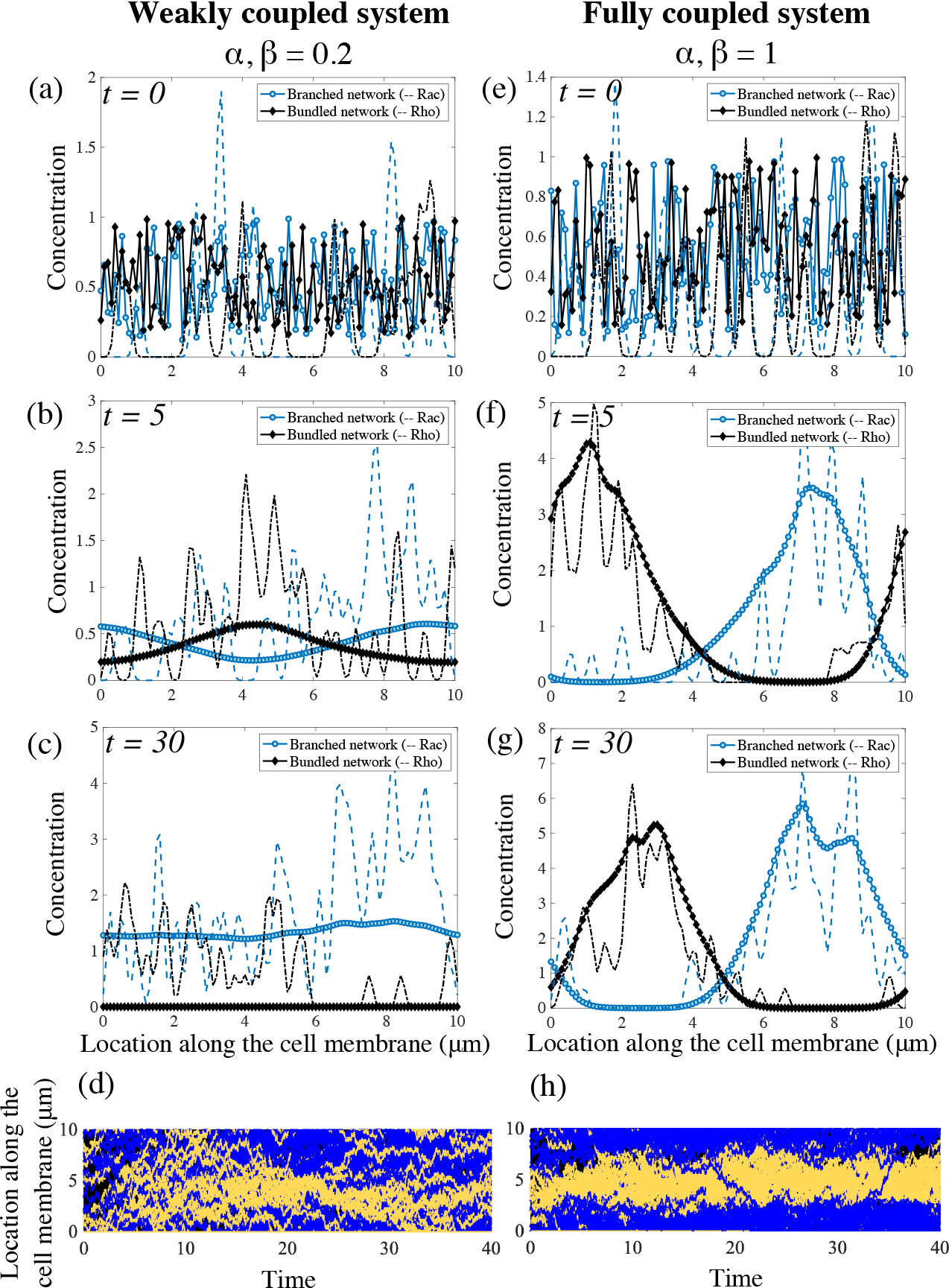
Emergence of polarity in the coupled mechanochemical system. (a–c) With weak feedback between the polarity molecules and the actin networks, the underlying chemicals do not segregate into a polar distribution but remain in a ‘patchy’ distribution around the cell membrane. In this particular simulation, the branched actin persists and eliminates the bundled actin meshwork in the system. (e–g) For strong coupling constants, both the mechanical and signaling system polarize. The polarity proteins completely segregate and each actin network separates into a peak of activity in complementary regions of the cell membrane. (d,h) Kymographs of the molecular locations of Rac and Rho around the cell edge. Rac/Rho trajectories are yellow/blue, respectively. The weak/strong coupling systems are (d)/(h), respectively.

Next, we considered a fully coupled system and observed that the polarity molecules segregate into two clusters on the membrane: a Rac-dominated front and a Rho-dominated rear (Fig. 3h). Simultaneously, the same spatial pattern is adopted by the actin networks: in the Rac-dominated region, branched actin network is assembled, while in the Rho-dominated patch bundled actomyosin is present (Fig. 3g). Initial patches of Rac enhance local recruitment of branched actin network while at the same time, in different regions Rho patches promote the formation of actomyosin bundles. This initial feedback-based recruitment leads to the formation of peaks in the concentration of each actin network, which ultimately increases the association and feedback rates in the signaling system. This transient behavior ultimately gives rise to a well-formed and stable peak in each actin species and a corresponding peak in its associated polarity molecule concentration. While these peaks are dynamic, their locations in space remains fixed given no external stimulus. Furthermore, perturbations of the spatial profile of the actin networks quickly return to equilibrium.

To characterize the parameter dependence on establishing polarity in the model, we performed simulations where all biochemical signaling and mechanical parameters are held constant but the coupling constants are varied. By simultaneously changing these two coupling strengths, we obtain contour plots of the polarization probability (Fig. 4a), defined as the fraction of the number of simulations with stable polar distributions of the actin networks and polarity molecules over the total number of runs. Based on Fig. 4, the system is more likely to polarize in the presence of higher coupling constants. Importantly, both feedback directions – from actin to Rho GTPases and from Rho GTPases to actin – are needed for the polarization.

**Figure 4:**
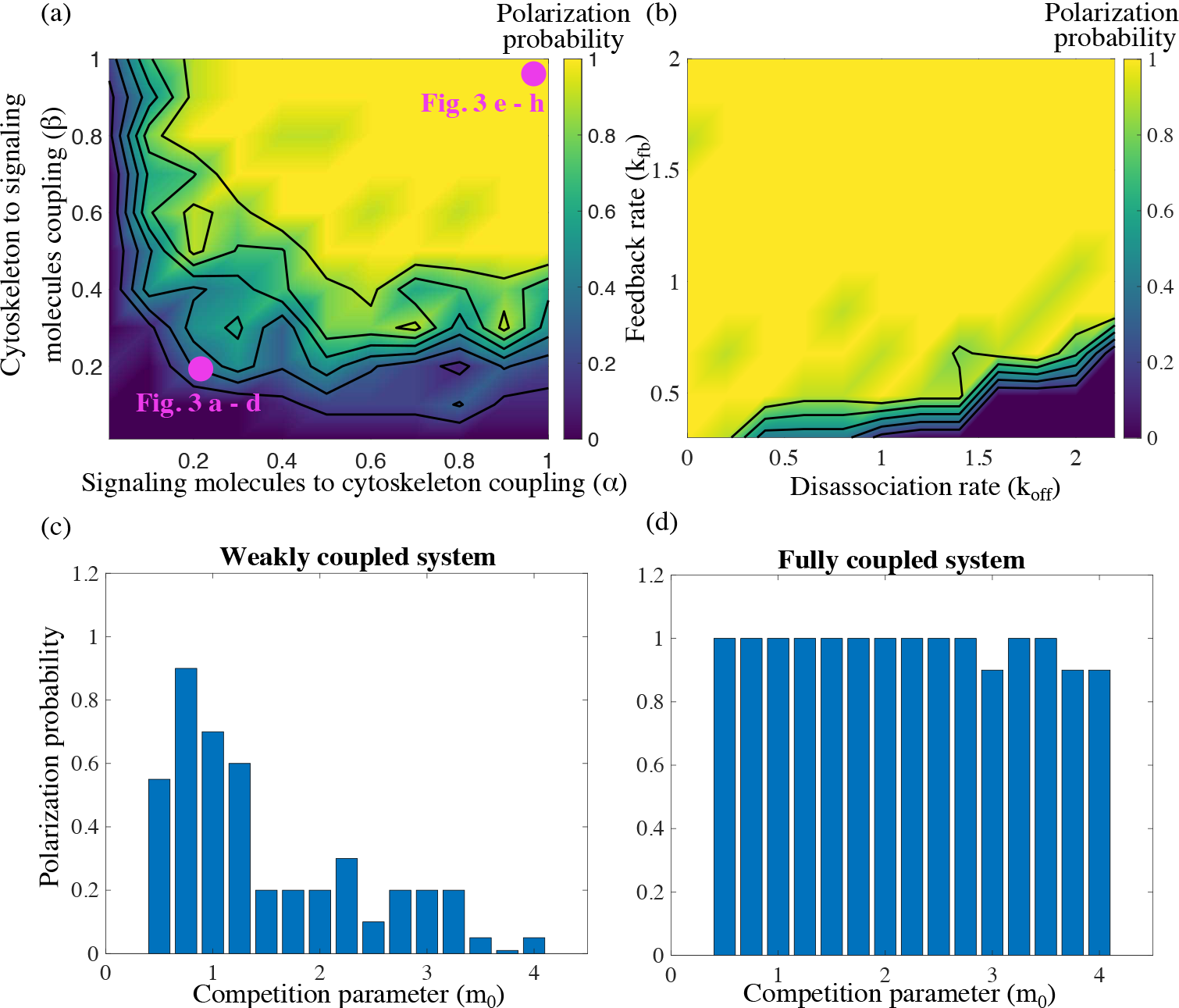
Stronger mutual coupling between the actin networks and the signaling system improves cell polarization. Ten simulations are considered for each set of parameters. Based on the outcome, a probability of a stable polarized solution is reported as the fraction of polarized solutions out of the total number of simulations for that specific choice of parameters. (a) Polarization probability is reported as a function of the two coupling parameters in the mechanochemical positive feedback loop, *α* and *β*. The parameters which were used to produce simulations shown in Fig. 3 are indicated. (b) Polarization probability as a function of two signaling kinetic rates: induced recruitment to the membrane rate *k*_fb_ and disassociation rate *k*_off_. (c–d) The influence of the actin competition parameter *m*_0_ on the polarization probability is reported for (c) a weakly coupled system, *α* = *β* = 0.2, and (d) a fully coupled system, *α* = *β* = 1.

### The effect of mechanical and biochemical rates on polarity establishment

We performed a series of simulations varying the competition parameter in the mechanical model, and obtained the corresponding polarization probability. For a fully coupled system, the competition parameter has little to no effect on the likeliness to establish polarity (Fig. 4d). In contrast, for a system with weaker coupling, the ability to establish a polar distribution of the actin networks is enhanced for lower competition parameters (Fig. 4c). In the absence of competition, the equations are well-known to exhibit coexistence with uniform spatial distribution [13]. We find that in the absence of competition, but coupled to the signaling module, both actin networks can coexist on the entire domain, but, due to the randomness of the signaling module, two actin densities exhibit arbitrarily positioned small-amplitude peaks. We also varied the relative initial amounts of branched and bundled actin networks and found that even in the case where 90% of the F-actin is initially assembled into actomyosin bundles, the cell was able to polarize with half of the actin in one type of network, and half in the other (Supplemental Material, Fig. S1a).

Next, we varied the kinetic rates in the biochemical signaling module. The mechanical parameters and coupling constants are held constant, while two chemical rates for both Rac and Rho were varied: the induced association rate, *k*_fb_, and the disassociation rate, *k*_off_. We obtained contour plots of the polarization probability (Fig. 4b). For the majority of the contour plot, the probability to polarize is largely unaffected by variations in these parameters. However, the model is sensitive to high disassociation rates and simultaneously, low enhanced recruitment rates (bottom right corner of Fig. 4b). In this situation of high disassociation and low feedback rates, there are too few polarity molecules on the cell membrane, hence the chemical system is not able to influence the mechanical module, and the system gravitates towards the stably ‘winner takes all’ state; one of the actin meshworks completely annihilates another.

The total number of polarity molecules in the system is also varied. Altschuler et al. [2] reported that for a large number of polarity molecules, the active molecules are spread over the membrane, and polarity is lost. By contrast, our coupled model does not exhibit the same response – the probability to polarize is largely unaffected by the total number of polarity molecules (Supplemental Material, Fig. S1b). Lastly, we report on the response to lowering the diffusion constant by an order of magnitude for a fully coupled system with baseline parameter values. We find that two actin density peaks do emerge, one corresponding to a protrusive zone with Rac and branched actin present, and a second peak with Rho and bundled actomyosin (Supplemental Material, Fig. S2). The width of these peaks is notably narrower than those reported in our other results due to the lower diffusivity. These peaks are also closer to each other. While the cell is, formally speaking, polarized, it does not have a well defined front and rear at opposite ends. Instead, the protrusive and retractive regions could be close to each other on the cell edge.

### Polarization direction is responsive to orientation of external signal

Thus far, our model simulations suggest that spontaneous polymerization in an arbitrary direction can arise from the coupling between the cytoskeletal mechanics and chemical kinetics. To determine whether our model exhibits sensitivity and adaptation to external signals, we simulated polarization in the presence of directional bias (Fig. 5 a–b). We assumed that the association/dissociation rates for Rac molecules vary along the cell edge, which is equivalent to a directional bias, as shown in Fig. 5a (dissociation rate varied oppositely to the association rate, as the spatial complement of the curve in Fig. 5a: the sum of the association and disassociation rates is constant). The kinetic rates for Rho molecules vary oppositely to those of Rac: where Rac rates were low, Rho rates were high and vice versa. We observed that a polarized state evolved from random initial conditions, with Rac and Rho peaks with the same orientation as the external bias (Fig. 5b). The actin networks peaks also co-localized with respective chemicals and the external bias (Fig. 5b).

**Figure 5:**
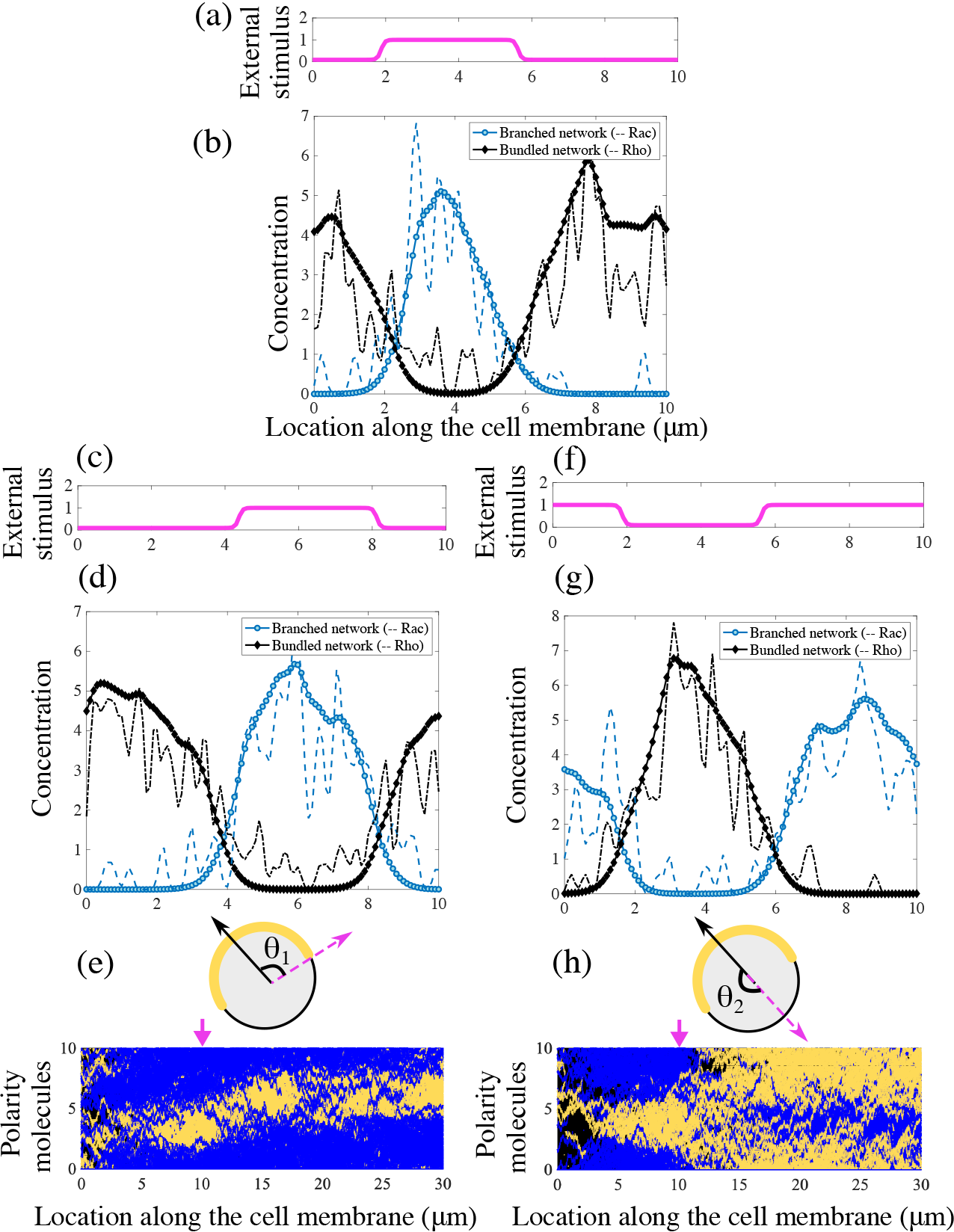
Polarization direction responds to changes in the external signal.. (a) An external stimulus imposes a directional bias on the kinetic rates of both polarity proteins Rac and Rho. Rac association and Rho disassociation rates have the profile around the cell edge shown in this plot. (b) With the biased chemical kinetics, polarized distributions of both signaling chemical concentrations and actin networks are oriented in the direction of this external stimulus. (c) After the polarized distributions shown in (b) evolve, the peak of the external stimulus is shifted by *θ*_1_ = 90 degrees. (d) In response to the shift of the external signal, the cell re-establishes polarity in the new direction. (e) Schematic of the change in external stimulus and resulting kymograph of the membrane-bound polarity proteins Rac/Rho (yellow/blue) along the cell membrane. (f) The peak of the external stimulus is now shifted by *θ*_2_ = 180 degrees, corresponding to a reversal of the incoming signal. (g) In response to the shift of the external stimulus, the cell re-establishes polarity in the opposite direction. (h) Schematic of the change in external stimulus and kymograph of the membrane-bound polarity proteins along the cell membrane in response to reversal of the external stimulus. Yellow indicates Rac chemicals, while blue indicates Rho chemicals. The arrows in the bottom plots indicate the temporal location of a new stimulus.

Next, we assessed how an already polarized cell responds to changes in the external signal direction. To achieve this, after the oriented polarized state established, we abruptly shifted the signal direction by 90 degrees, Fig. 5 c, e). After a transient reorganization of the signaling chemicals, the actin networks rearrange as well and the polarization direction turns to the new direction (Fig. 5 d, e). If this was a motile cell, it would execute a smooth turn: at no time the polarization was lost, just its orientation changed smoothly. Lastly, we reversed the direction of the external signal by 180 degrees (Fig. 5 f, h), and observed that the cell re-polarized in the new direction (Fig. 5 g, h). Note that in this case, polarization was momentarily lost but emerged in a new direction.

### The model is robust to changes in initial and boundary conditions

To investigate the sensitivity to initial conditions, we explore the models response to different initial distributions of the actin networks. A feature of robust polarity establishment is the ability to coalesce multiple protrusions, or peaks in the branched actin concentration, into a single protrusive front. Thus, we considered an initiation of the system where branched actin concentration has three evenly spaced peaks, bundled actin density has one square peak, but active signaling molecules concentrations are randomly distributed along the plasma membrane (Fig. 6a). All ten instances of this simulation setup showed the formation of a single protruding front (Fig. 6a). The location along the cell membrane of this front will vary based on the initiation.

**Figure 6:**
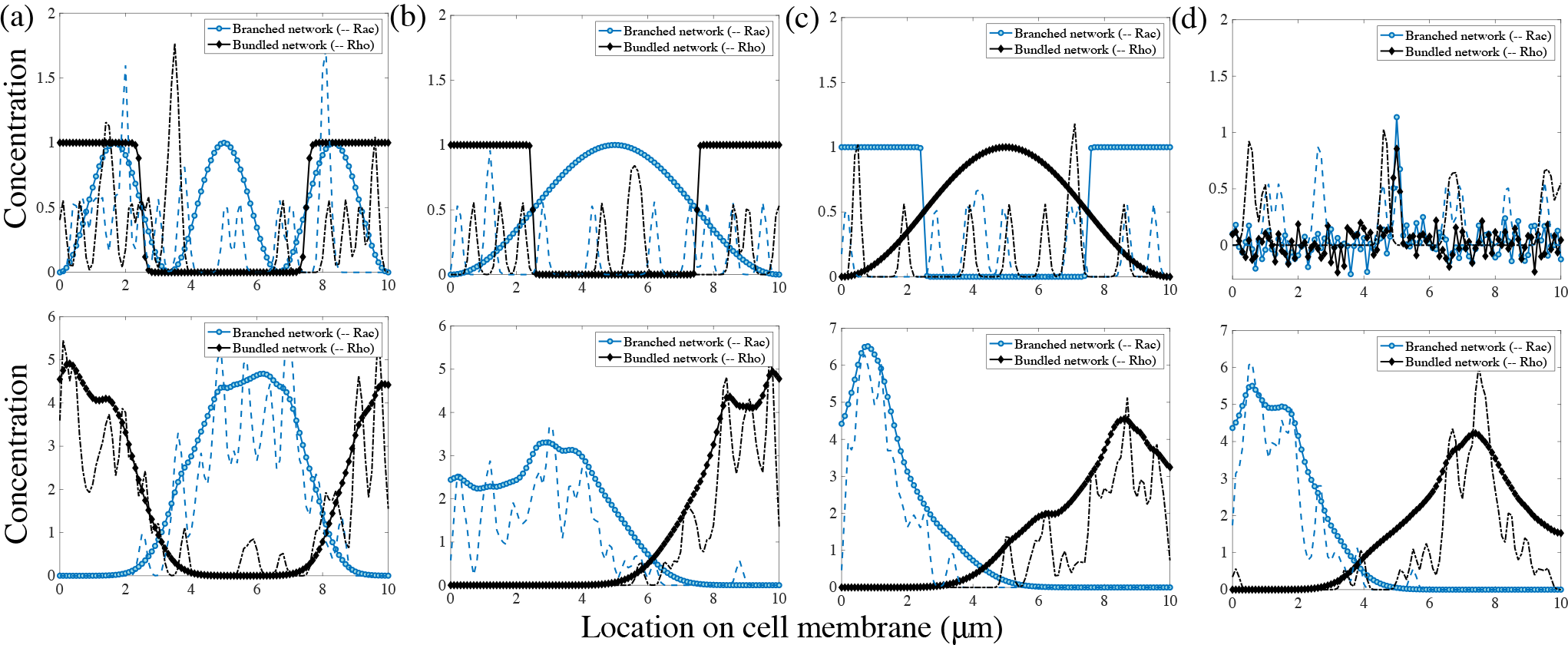
Polarity establishment is robust to variations of the initial distributions of the cytoskeletal meshworks and of the boundary conditions. Initial (top row) and evolved (bottom row) distributions of the actin networks and polarity molecules. In all initial conditions, the Rac and Rho molecular distributions are random. (a) Periodic boundary conditions are used representing the circular cell edge. Initially, there is one peak of the bundled network density but three peaks of the branched network density, two of which co-localized with the bundled network. (b)–(d) The model on the 1D segment with no-flux boundary conditions representing the anterior-posterior cross-section of an elongated cell. (b) Initially, the ends of the cell are contractile, while the middle is protrusive. (c) Initially, the cell ends are protrusive, while the middle is contractile. (d) Initially, both protrusive and contractile networks are concentrated at the cell center. In all cases, the same polarized state with segregated protrusive and contractile networks evolve. Notably, for the segment-like cell, the networks segregate to the opposite ends of the cell making the cell robustly motile.

Next, the model geometry is changed: instead of the circular edge of a disc-like cell, we considered the anterior-posterior cross-section of an elongated cell. The model remains 1D, but the boundary condition becomes no-flux, as actin, myosin and signaling molecules do not enter or exit the cell. All previously described results in this case qualitatively match those obtained by solving the same set of equations on the 1D segment with periodic boundary conditions (see Supplemental Material, Figs. S3–S4). We show that three possible different initial spatial distributions of actin networks all evolve to a stable polarized state with protrusive network at one end of the 1D cell, and contractile network at the other end (Fig. 6 b–d).

## Discussion

The initiation of cell migration involves a complex web of signaling and cytoskeletal modules. While in certain cellular systems polarity can arise from only signaling or mechanical pathways, many cells rely on the interplay between both to robustly break symmetry to initiate locomotion. We have presented a mechanochemical model for cell polarization based on two minimal sub-models, one describing signaling molecular dynamics, another mechanics of cytoskeletal networks. The first sub-model is for two types of signaling molecules, like Rac and Rho, which can diffuse on the cell membrane, dissociate from the membrane or move from the cytoplasm to the membrane both spontaneously, or can be recruited to the membrane by other molecules of the same type. This induced recruitment is the simplest autocatalytic feedback and corresponds to the only nonlinear (quadratic) term in the mathematical model. One species of molecules with these dynamics aggregates into a finite number of clusters [2]. Steric interaction is the only relation between two different types of molecules in the model. We show that such interaction leads to the emergence of multiple interspersed clusters across the cell. This signaling system does not polarize the cell globally: different type molecules locally segregate, but the clusters of each kind do not aggregate in respective halves of the cell, as required for stable cell polarization.

The second sub-model is for two types of dynamic actin networks at the cell edge, branched protruding meshwork and actin-myosin contractile bundled network. These networks slowly and randomly spread around the cell edge, due to physical movements and treadmilling of actin filaments, and turn over while maintaining certain equilibrium density. The nontrivial interaction between these networks is competition, such that the local density of one tends to diminish the density of another. This interaction stems from both mechanical effects and from competition for molecular resources. Mathematically, it is described by the simplest quadratic nonlinear term, which makes this sub-model equivalent to the Lotka-Volterra equations for two competing populations [13]. It was shown in [32] that this competition between two actin networks is an important part of the spontaneous polarization process, but without cell movement, the model is not able to polarize the cell, as one network will always win.

Thus, neither model is capable of producing cell polarization on its own. Here, we demonstrated that the simplest coupling between the chemical and mechanical models local, linear positive feedback loops between signaling molecules and actin cytoskeletal networks is sufficient for spontaneous polarity. The model works if the strength of the mechanochemical coupling is above a certain threshold. Qualitative mechanism suggested by the model is simple: branched/bundled actin networks support recruitment of Rac/Rho to the membrane, respectively, so Rac and Rho tend to segregate into separate parts of the cell. In turn, neither network can now invade anothers territory because Rac/Rho enhance branched/bundled networks, respectively; for example, when the bundled network tries to encroach into the part of the cell occupied by the branched actin, Rac co-localizing with branched actin gives its advantage over bundled actin and prevents the invasion.

The competition between the actin networks has to be strong enough for stable polarization, so that the networks do not coexist locally, but not too strong to overcome the effect of the mechanochemical coupling and enable one network to win the cell from the other. Effective diffusion also has to be neither too slow, nor too fast for the model to work. If the diffusion coefficients in the model are too small, then the different networks with their supporting signal molecules localize into peaks that are too narrow and arbitrarily positioned. If the diffusion coefficients are too large, no pattern forms. However, overall the model is robust: a few fold variations of any other parameters do not change the stable separation of branched actin and Rac in one half of the cell, and of bundled actomyosin and Rho to another half. The polarized pattern is not sensitive to the initial or boundary conditions. It is very likely that the model performs in 2D and 3D as well as in 1D. Interestingly, the model works even if the total number of signaling molecules becomes too large, while in this regime, the signaling sub-model without mechanochemical coupling fails to produce clustering [2]. Likely, this means that a fully continuous variant of the mechanochemical model also supports stable polarization.

Our simulations showed that the model cell can polarize spontaneously, without any signal from the environment, but the direction of polarization adapts to an external signal. For example, if the kinetics of the polarity molecules are biased in a certain direction by an external signal, then the cell polarizes in that specific direction. If the cell is already polarized, and the signal is applied in a different direction, this causes a reorientation of the entire mechanochemical polarity machinery in the new direction. If the signals direction is close to that of the cell orientation, then the mechanochemical pattern turns smoothly, otherwise, the cell first momentarily looses polarity, and then re-polarizes in the new direction. This response is similar to experimental observations: motile cells execute a smooth turn when the external signal is applied normally to the direction of locomotion [1], but re-polarize when the signal is opposite to the movement direction [59].

The first conceptual biological implication of our model is a mathematical demonstration that a signaling module coupled to a cytoskeletal module leads to robust spontaneous polarization and reorientation in the presence of incoming signals; this occurs despite the fact that each module separately can segregate chemical/networks in space but cannot stably polarize the cell. Second, there are a number of solely chemical and solely mechanical polarization models, but for these to work, significantly nonlinear terms, supporting cooperative or switch-like behavior, are required. We show that a linear local coupling of two minimally nonlinear models (only quadratic nonlinearity in actin growth) can achieve robust cell symmetry breaking.

Our model conceptually relates to a number of cell types. The first example is neutrophils, which spontaneously polarize and require intimate involvement of actin dynamics in the polarization process. These cells, when polarized, follow changes in external signals, and a qualitative cartoon model, extremely similar in spirit to our model, was proposed [72]. More recent experimental findings also support the coupled action of actin networks and Rho GTPases in spontaneous neutrophil polarization [9, 20]. Similarly, polarization of fibroblasts, macrophages, and astrocytes is certainly mechanochemical, not purely chemical process, with cytoskeletal dynamics intimately intertwined with RhoGTPase signaling in the symmetry breaking process prior to migration (reviewed in [15]).

We do not claim that our model can predict the biological details of polarity for these cell types. In general, polarization involves not only actin, but also microtubules, as well as more complex cytoskeletal networks, such as stress fibers, multiple pseudopods, and adhesion complexes. Signaling circuits other than Rho GTPases could also be implicated in cell polarization. Even limited to just four molecular players – Rac, Rho and two actin structures – the model is extremely simplified and is conceptual rather than detailed. Or model posits one of the simplest quantitative frameworks for understanding a possible mechanism for spontaneous mechanochemical cell polarization.

It is possible for our model to rely on other forms of feedback between the biochemical and mechanical circuits. For example, negative, instead of positive, feedback between Rac and branched actin, and Rho and actomyosin, respectively, could do the job [70, 72]. We also limited the dynamics of the model to the local chemical and mechanical processes, but global mechanical effects, for example, membrane tension, could play an important role in polarization of some cell types [25]. Another paradigm for mechanochemical polarization requires transport of chemicals in the signaling framework. The key for such models is that myosin-driven flow assists the polarization of signaling proteins by mechanically triggering the formation of a stable asymmetric chemical distribution [18, 35, 62]. Our model is simpler because it does not have directional movement – either in the form of a flow as in these models, or in the form of whole cell movement as in [32]. More detailed and complex models have included the cell-surface adhesion dynamics as a mechanical component in the biochemical polarization pathway [48]. Finally, one of the models of gradient sensing [17] is based on an idea similar to our model: when signaling dynamics are such that clusters of polarity molecules appear on the cell membrane, the clusters’ location can be biased to one side of the cell by an external signal. Our model adds a novel and simple potential polarization mechanism to these theoretical paradigms.

## Author Contributions

C.C. and A.M. designed the project and developed the models. C.C. carried out the hybrid stochastic-deterministic simulations and analyzed the data. C.C. and A.M. wrote the manuscript.

## Acknowledgments

This work was supported in part by US Army Research Office grant W911NF-17-1-0417 to C.C. and A.M.

## Supplementary Material

### S.1 A general description of the model

The model considers the dynamics of two actin networks competing for molecular resources and coupled to the dynamics of active, membrane-bound Rac and Rho molecules. In the model, the dynamics is localized to the periphery of the the disc-shaped cell adhering to a substrate, and so the molecular densities are localized to the circle of circumferential length *L*.

#### Actin dynamics

The cytoskeletal model is a competition of two distinct actin networks with the following dynamics:

1. *Autocatalytic growth*: The net growth rate of each network is proportional to local network density. This assumption is based on the processes of polymerization of existent actin filaments and of nucleation of nascent filaments by proteins binding to the existent filaments, so that the net growth becomes proportional to the existent density.
2. *Limited growth*: At high density, growth is limited due to lack of availability of molecular resources. In the case of the bundled actin network, growth could be limited due to depletion of the myosin-II motors or actin monomers, while the branched actin network growth could be limited by availability of Arp2/3 branching complexes or globular actin monomers.
3. *Competition for molecular resources*: Both networks compete for a limited cytoplasmic pool of molecular resources, such as G-actin monomers, Arp2/3 complexes, formins or myosin.
4. *Diffusive-driven redistribution of the networks along the cell boundary*: We assume an effective diffusive movement of actin along the cell edge due to lateral shifts of the actin density due to filament growth and/or to physical sliding of filaments along the cell edge pulled by molecular motors.

Mathematically, based on these assumed dynamics one arrives at the following set of non-dimensionalized PDEs [32]:

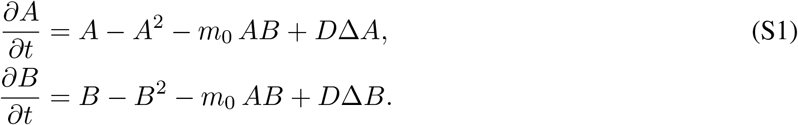

Here *A*(*s*, *t*) denotes the branched actin network density and *B*(*s*, *t*) represents the bundled actin network density along the cell boundary parameterized by the arc length *s*. Densities of both actin networks are defined on the periodic cell boundary. *m*_0_ is the non-dimensional competition parameter, and *D* is the non-dimensional diffusion coefficient.

#### Signaling molecule dynamics

We focus on the mutually exclusive interactions between Rac1 and RhoA on the plasma membrane. Following the rationale of the stochastic model proposed by Altschuler et al. [1], we assume five different kinds of molecular events:

1. *Spontaneous association to the membrane*: GTP-bound RhoGTPase proteins undergo a conformational change and transition to an active membrane-bound state. We model this by an association of a respective molecule from the cytosol to a random location on the membrane at a rate of *k*_on_.
2. *Spontaneous disassociation from the membrane*: GAP proteins regulate the transition of active, membrane-bound RhoGTPases into an inactive, cytosolic state. This event is modeled through the removal of an active molecule from the membrane at a rate of *k*_off_.
3. *Enhanced membrane association through activators*: Local positive feedback loops are thought to play a role in sustaining nascent Rac/Rho sites on the plasma membrane [2–5]. To model these feedback loops we assume that a membrane-bound (active) molecule of either type (Rac or Rho) can indirectly activate and recruit a molecule of the same type to its vicinity. The rate at which one molecule recruits from the cytosol is proportional to the fraction of molecules which are still in the cytosol with a proportionality constant of *k*_fb_.
4. *Diffusion on the membrane*: Each molecule on the membrane undergoes a Brownian motion with diffusion coefficient *D*.
5. *Steric interaction*: In the association, recruitment, and diffusive processes, Rac and Rho proteins cannot occupy the same location in space at a given time. This assumption is based on the reported mutual antagonistic interactions between RhoGTPases [6–11].

In the stochastic approach to the molecular dynamics, the number of Rac molecules on the membrane *n^Rac^*(*t*) evolves by a continuous time Markov process. In the case of position-independent dynamics (in the absence of the feedback from actin to Rac), the probability of *n^Rac^* molecules present on the membrane evolves according to the following master equation:

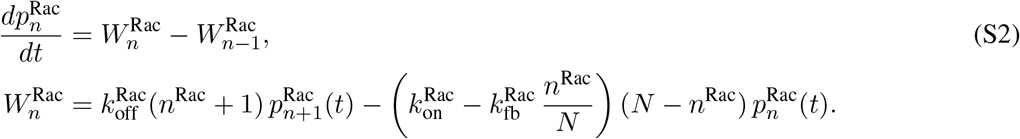

Here *N* is the total number of Rac molecules in the cell. We write similar equations for the number of Rho particles on the membrane at a given time *n^Rho^*(*t*):

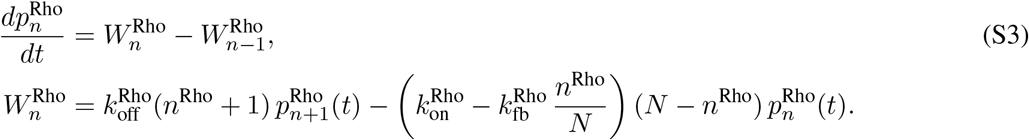

The molecules dissociate from the membrane with a constant rate, independent of their location. The molecules associate with the membrane with the rate dependent on the number of molecules already on the membrane. They associate with certain probability to a random location on the membrane; otherwise, they associate with another, randomly chosen, molecule already on the membrane. This probability and the numerical procedure are described below. In the presence of the mechanochemical coupling, the rates of association and dissociation depend on position on the cell edge, and the equations are modified as discussed below.

#### Mechanochemical coupling

For the mutual coupling between actin cytoskeleton and polarity molecules, we assume that there is a local feedback loop with a linear dependence on relative concentrations. The chemical rates in the signaling kinetics are no longer constant but rather dependent on the local concentration of each respective actin network which evolves in both space and time. We assume that Rac and the branched actin network engage in a positive feedback loop and similarly so do Rho and the bundled actomyosin mesh by modifying the kinetic rates of Rac and Rho as follows:

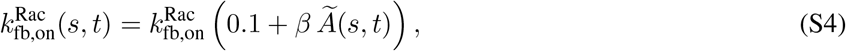

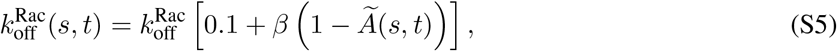

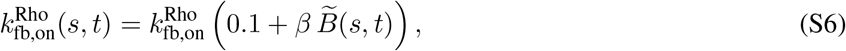

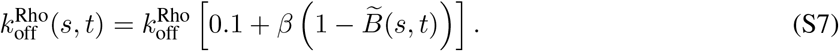

The strength of the coupling from actin to the polarity model is denoted by the constant of proportionality *β*. The tilde symbol indicates that these concentrations of actin networks have been normalized by their maximum value so that at a given time concentration of actin varies spatially between 0 and 1. On the reverse, the growth rate of each actin network is now an evolving parameter that depends linearly on the local amount of active or membrane-bound polarity proteins:

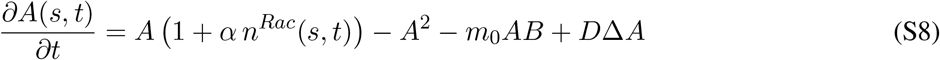

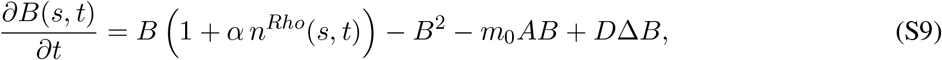

where *α* represents the strength of the coupling from the polarity molecules to the cytoskeleton.

**Table S1:** Definition and values of parameters for the hybrid mechanochemical polarity model.

| Parameter | Value | Description | Reported value | Reference |
| --- | --- | --- | --- | --- |
| $L$ | $10 \mu\text{m}$ | Length of cell | $\sim 5 - 20 \mu\text{m}$ | [12] |
| $D$ | $0.5 \mu\text{m}^2/\text{sec}$ | Effective diffusion of molecules on membrane | $0.02 - 0.5 \mu\text{m}^2/\text{sec}$ | [1, 13–15] |
| $m_0$ | 2 | Competition or bundling term | | Chosen from compartment model simulations in order to give rise to quasi-stable polar solutions (varied). |
| $N$ | 200 | Rho GTPase molecules in cell (conserved) | | Chosen from Altschuler et al. [1] in order to give rise to patches (varied). |
| $k_{\text{on}}$ | $0.001/\text{sec}$ | Association rate for Rho GTPases | $1.67 \times 10^{-5}/\text{sec}$-$0.027/\text{sec}$ | [1, 15] |
| $k_{\text{fb}}$ | $1/\text{sec}$ | Autocatalytic activation rate for RhoGTPases | $0.1667/\text{sec}$ | [1] |
| $k_{\text{off}}$ | $0.9/\text{sec}$ | Disassociation rate for RhoGTPases | $1/\text{sec}$, $0.15/\text{sec}$, $0.02/\text{sec}$ | [1, 14, 16–18] |
| $h_{\text{eq}}$ | 0.1 | Fraction of membrane-bound RhoGTPases | 2-10% | [1, 15] |
| $\epsilon$ | $0.01 \mu\text{m}^2$ | Variance of Gaussian function used sampling Rac/Rho concentrations | | |

#### Numerical simulations

The theoretical approach provided above could describe the actin networks concentration and polarity molecules on a one-dimensional curve or a two-dimensional surface of the plasma membrane. The numerical simulations carried out here were on one-dimensional circles for ease of visualization, but we believe the results here could be reproduced in higher dimensions on arbitrary geometries. To simulate the dynamics of cell polarization, the computational domain representing concentrations in the plasma membrane and a thin volume of cytoplasm adjacent to the membrane is discretized using 101 points with an averaged spatial discretization of ∆*s* = 0.1 *μ*m. The temporal discretization is ∆*t* = 0.01 sec and simulations are run to 30-100 seconds. The codes are written and solved in Matlab. Model parameters along with justifications for the choice of values are provided in supplementary material, Table S1. We perform simulations using the baseline parameter values listed in Table S1 unless otherwise indicated.

The actin dynamics PDEs in Eq. S1 are solved on a circular domain using Crank-Nicolson finite difference numerical method with periodic boundary conditions. The actin networks are randomly distributed initially with equal relative concentrations between branched and bundled networks.

A Gillespie algorithm is used for the stochastic equations for the polarity molecules. The equations in Eqs. S2 and S3 completely determine the random evolution of the membrane content for Rac/Rho particles, and the resulting Markov process generates a sequence of random times *t*_1_ < *t*_2_ < *t*_3_ < … at which either the number of Rac or Rho molecules on the membrane jumps by +1 or −1. In between these time points, the number of either molecules on the membrane is constant. In between the jumps, the molecules with locations 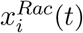 and 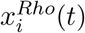, where *i* is the index of the specific molecule, undergo Brownian motion on the membrane with diffusion coefficient 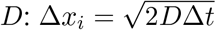. Since we enforce segregation of Rac and Rho, collisions between Rac and Rho molecules in the diffusive process may occur. We resolve collision events by not allowing either molecule to move into the space (interval of width ∆*x* around a given molecule) that would result in overlap. Other more sophisticated collision resolution methods could have been employed, but for simplicity we chose this minimal dynamic.

In the Markov process, if an association with the membrane occurs, spontaneously or by the feedback from molecules already on the membrane, we augment the appropriate list of molecules and choose a location for this new molecule. The location of this new molecule depends on whether the association event is spontaneous or by the feedback. Following the analysis in Altschuler et al. [1], a spontaneous association event occurs with probability *k*_on_/ (*k*_on_ + *k*_fb_ · (*N* − *n*(*t_i_*))), in which case the location of the new molecule is randomly chosen such that it does not spatially overlap with any existing molecule of a different type (the molecule arriving on the membrane cannot be inside the intervals of width ∆*x* around any existing molecule of a different type). Here, *n* can refer to either the number of Rac or Rho particles since the rules that govern their dynamics are the same. Otherwise, a feedback-driven association occurred and the location of the new molecule is chosen to be within the interval ∆*x* of a randomly chosen already membrane-bound molecule of the same type.

In the case there is the feedback from the actin densities to the Rac/Rho system, the association and dis-association rates start to be time- and position-dependent. Then, in the equations for the total numbers of the molecules on the membrane, the association (both spontaneous and feedback-driven) and disassociation rates at each given time are the spatial averages of the actin density-dependent rates given by equations Eqs. S4–S7. When a dissociation event occurs, the molecule to be dissociated is not picked from all such molecules on the membrane with equal probability, but with a probability proportional to the local dissociation rate given by Eqs. S5,S7. Similarly, when a feedback-driven association event occurs, the new molecule is associated with any molecule already on the membrane not with equal probability, but with a probability proportional to the local feedback-driven association rate given by Eqs. S4,S6. The actin density in formulas S4–S7, which is known on the discretized grid, is extrapolated to the actual positions of the molecules on the membrane using standard linear extrapolation.

Lastly, to complete the numerical algorithm, we must specify how we initialize the system. At the start of the simulation, 10 % of the total conserved number of each particle type are chosen to be in their active, membrane-bound state. Their locations are randomly picked along the cell membrane in a way to ensure that Rac and Rho particles do not overlap.

**Figure S1:**
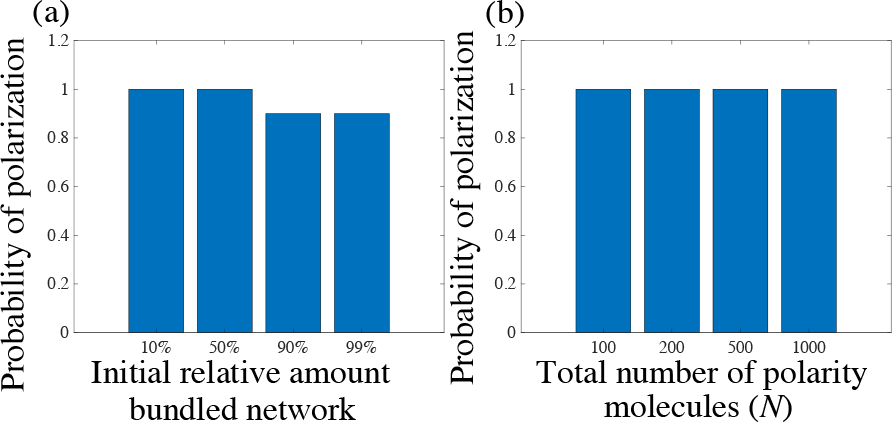
Model sensitivity to variations in total number of polarity molecules and relative initial concentrations of actin. (a) Probability of polarization as a function of the total number of polarity molecules, *N*. (b) Probability of polarization as a function of the initial amount of bundled actomyosin network relative to branched network.

**Figure S2:**
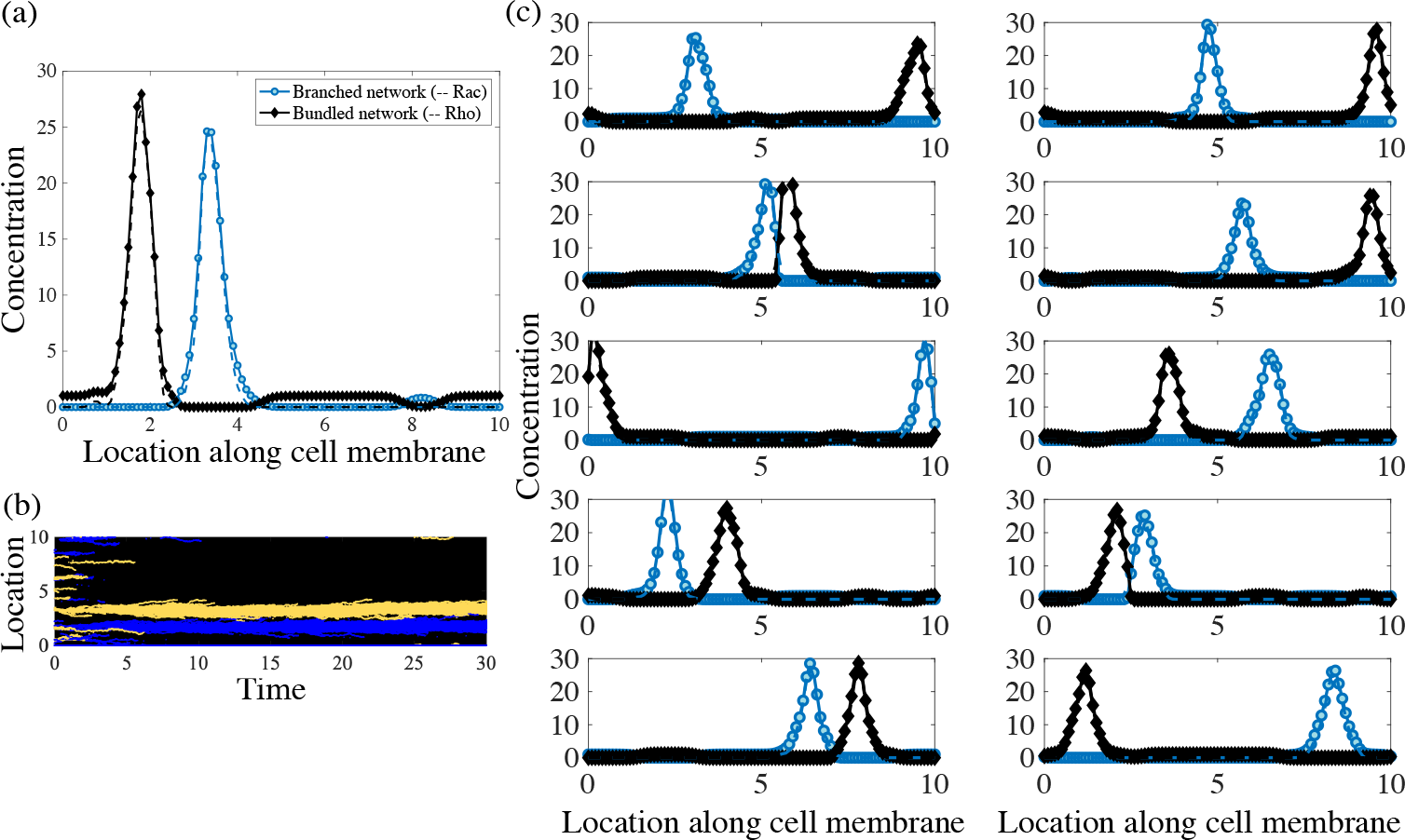
Model response with a lower diffusion constant of *D* = 0.01 *μ*m^2^/sec for both actin cytoskeleton and Rac/Rho systems. (a) One realization of the resulting distributions of both signaling chemical concentrations and actin networks. (b) The corresponding kymograph of the time-evolution of the active, membrane-bound polarity proteins. (c) Ten simulation results with a diffusion constant of *D* = 0.01 *μ*m^2^/sec. All other parameters including the coupling constants are held at their baseline values reported in Table S1.

### S.2 Model simulations with no-flux boundary conditions

We performed a series of simulations of the model for a one-dimensional anterior-posterior slice along the long axis of the cell in order to capture the dynamics of polarization in the anterior-posterior direction of elongated cells. We solved the system of coupled hybrid stochastic-deterministic equations (S1–9) with no flux boundary conditions to enforce the conservation of molecular numbers in this new geometry. Overall, the results of simulations qualitatively agree with the observations reported in the paper for periodic boundary conditions (Figs. S1 and S2).

**Figure S3:**
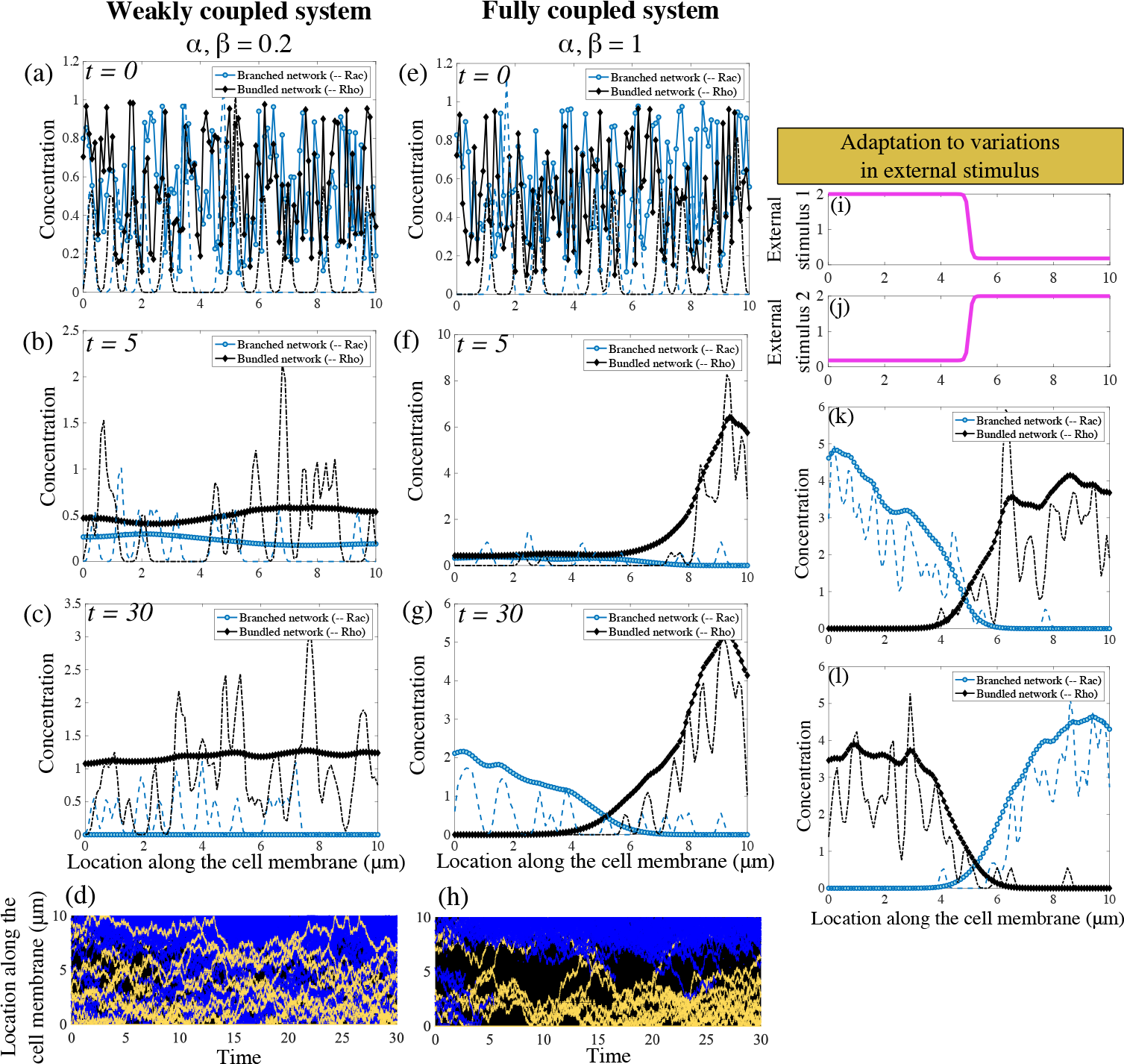
Polarized distributions are stable solutions of the mechanochemical hybrid model with no flux boundary conditions for large enough coupling constants. (a)-(d) With weak feedback coupling between polarity molecules and actin network, patchy initial conditions can result in non-polarized cells. (e)-(h) As the coupling constants are increased, both the mechanical and signaling systems polarize. (i)-(l) The model displays sensitivity to new incoming signals and both the cytoskeleton and signaling modules undergo rearrangement in the presence of a new external stimulus.

**Figure S4:**
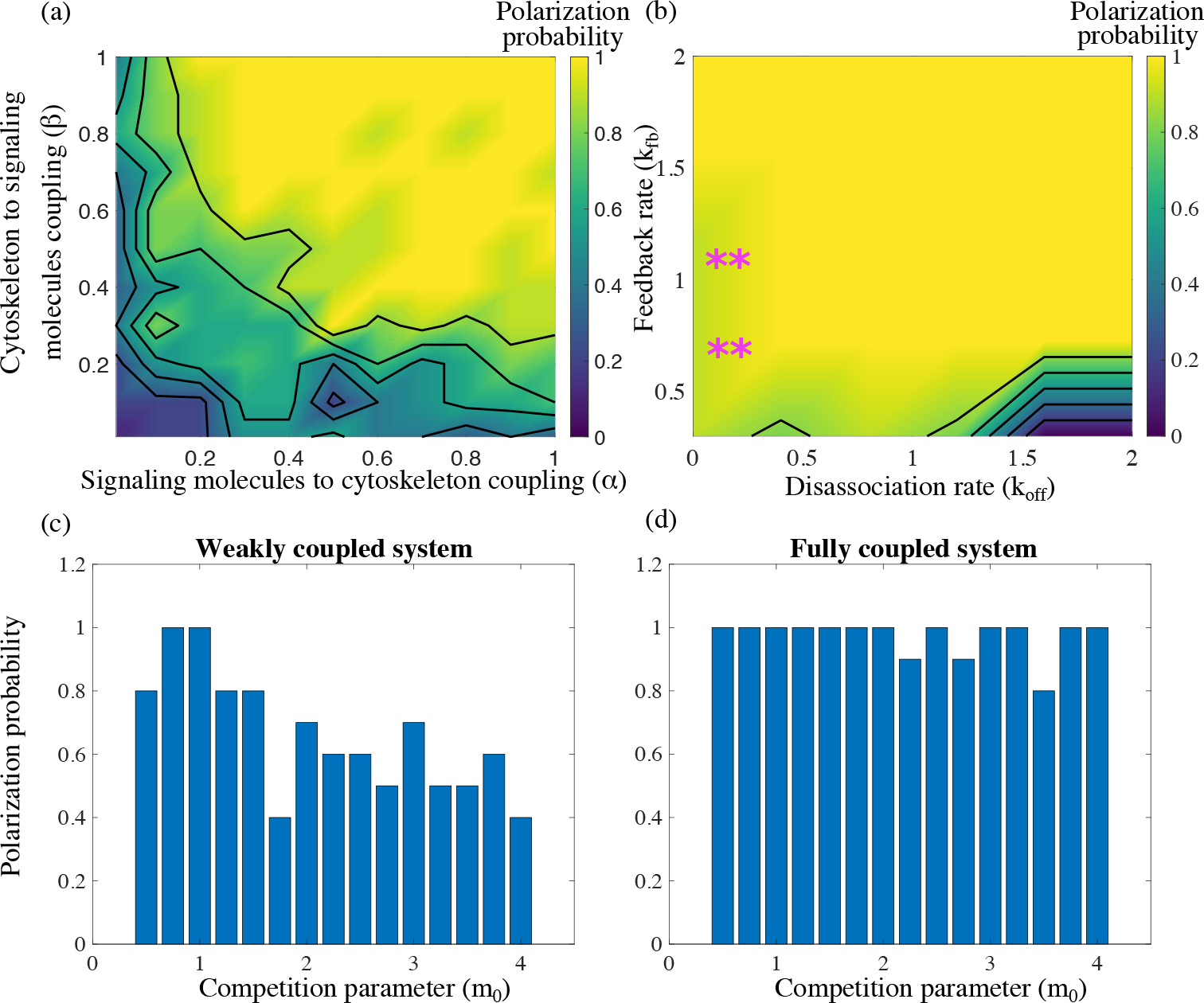
Sensitivity of the mechanochemical hybrid model with no flux boundary conditions. Ten simulations were done for each se t of parameters. Based on the outcome, a probability of a stable polarized solution is reported as the fraction of polarized solutions out of the total number of simulations for that specific choice of parameters. (a) Polarization probability is reported as a function of the two parameters, *α* and *β*, in the mechanochemical positive feedback loop. (b) Polarization probability is also reported as a function of the two signaling kinetic rates: feedback-driven association with the membrane *k*_fb_ and membrane disassociation *k*_off_. The asterisk (^*∗∗*^) indicates the existence of at least one simulation resulting in two stable peaks in either branched or bundled actin network. Such events were counted as failure to polarize. (c–d) Lastly, the dependence of the polarization probability on the competition parameter, *m*_0_, is reported for (c) a weakly coupled system, *α* = *β* = 0.2, and (d) a fully coupled system, *α* = *β* = 1.

## References

[1] Allen, G., Lee, K., Barnhart, E., Tsuchida, M., Wilson, C., Gutierrez, E., Groisman, A., Mogilner, A., and Theriot, J. (2019). Cell mechanics at the rear act to steer the direction of cell migration. bioRxiv, 443408.

[2] Altschuler, S., Angenent, S., Wang, Y., and Wu, L. (2008). On the spontaneous emergence of cell polarity. Nature, 454:886–889.

[3] Assemat, E., Bazellieres, E., Pallesi-Pocachard, E., le Bivic, A., and Massey-Harroche, D. (2008). Polarity complex proteins. Biochim. Biophys. Acta, 1778:614–630.

[4] Barnhart, E., Lee, K.-C., Allen, G., Theoriot, J., and Mogilner, A. (2015). Balance between cell-substrate adhesion and myosin contraction determines the frequency of motility initiation in fish keratocytes. Proc. Natl. Acad. Sci. USA, 112:5045–5050.

[5] Bois, J., Julicher, F., and Grill, S. (2011). Pattern formation in active fluids. Phys. Rev. Lett., 106:028103.

[6] Burridge, K. and Wennerberg, K. (2004). Rho and Rac take center stage. Cell, 116:167–179.

[7] Byrne, K., Monsefi, N., Dawson, J., Degasperi, A., Bukowski-Wills, J., Volinsky, N., Dobrzyski, M., Birtwistle, M., Tsyganov, M., Kiyatkin, A., Kida, K., Finch, A., Carragher, N., Kolch, W., Nguyen, L., von Kriegsheim, A., and Kholodenko, B. (2016). Bistability in the Rac1, PAK, and.RhoA signaling network drives actin cytoskeleton dynamics and cell motility switches. Cell Syst., 2:38–48.

[8] Cosner, C. and Lazer, A. (1984). Stable coexistence states in the Volterra-Lotka competition model with diffusion. SIAM J. Applied Math., 44:1112–1132.

[9] Dandekar, S. N., Park, J. S., Peng, G. E., Onuffer, J. J., Lim, W. A., and Weiner, O. D. (2013). Actin dynamics rapidly reset chemoattractant receptor sensitivity following adaptation in neutrophils. Philos. Trans. R. Soc. London B: Biol. Sci., 368(1629):20130008.

[10] Das, S., Yin, T., Yang, Q., Zhang, J., Wu, Y. I., and Yu, J. (2015). Single-molecule tracking of small GTPase Rac1 uncovers spatial regulation of membrane translocation and mechanism for polarized signaling. Proc. Nat. Acad. Sci. USA, 112:E267–E276.

[11] Dawes, A. and Munro, E. (2011). PAR-3 oligomerization may provide an actin-independent mechanism to maintain distinct PAR protein domains in the early *Caenorhabditis elegans* embryo. Biophys. J., 101:1412–1422.

[12] Dayel, M., Akin, O., Landeryou, M., Risca, V., Mogilner, A., and Mullins, R. (2009). In silico reconstitution of actin-based symmetry breaking and motility. PLoS Biol., 7:e1000201.

[13] Edelstein-Keshet, L. (1988). Mathematical Models in Biology. Random House, New York.

[14] Edelstein-Keshet, L., Holmes, W., Zajac, M., and Dutot, M. (2013). From simple to detailed models for cell polarization. Philos. Trans. R. Soc. London B: Biol. Sci., 368:20130003.

[15] Etienne-Manneville, S. (2006). In vitro assay of primary astrocyte migration as a tool to study Rho GTPase function in cell polarization. Methods Enzymol., 406:565–578.

[16] Falkenberg, C. and Loew, L. (2013). Computational analysis of Rho GTPase cycling. PLoS Comput. Biol., 9:e1002831.

[17] Gamba, A., Kolokolov, I., Lebedev, V., and Orentzi, G. (2007). Patch coalescence as a mechanism for eukaryotic directional sensing. Phys. Rev. Lett., 99:158101.

[18] Goehring, N., Trong, P., Bois, J., Chowdhury, D., Nicola, E., Hyman, A., and Grill, S. (2011). Polarization of PAR proteins by advective triggering of a pattern-forming system. Science, 334:11371141.

[19] Goryachev, A. and Leda, M. (2017). Many roads to symmetry breaking: molecular mechanisms and theoretical models of yeast cell polarity. Mol. Biol. Cell, 28:370–380.

[20] Graziano, B., Town, J., Nagy, T., Fonari, M., Peni, S., Igli, A., Kralj-Igli, V., Gov, N., Diz-Muoz, A., and Weiner, O. (2018). Cell-extrinsic mechanical forces restore neutrophil polarization in the absence of branched actin assembly. bioRxiv, 457119.

[21] Haglund, C. and Welch, M. (2011). Pathogens and polymers: microbe-host interactions illuminate the cytoskeleton. J. Cell Biol., 195:7–17.

[22] Hall, A. (1998). Rho GTPases and the actin cytoskeleton. Science, 279:509514.

[23] Heasman, S. and Ridley, A. (2008). Mammalian Rho GTPases: new insights into their functions from in vivo studies. Nat. Rev. Mol. Cell Biol., 9:690–701.

[24] Higgs, H. and Pollard, T. (2001). Regulation of actin filament network formation through Arp2/3 complex: activation by a diverse array of proteins. Annu. Rev. Biochem., 70:649676.

[25] Houk, A., Jilkine, A., Mejean, C., Boltyanskiy, R., Dufresne, E., Angenent, S., Altschuler, S., Wu, L., and Weiner, O. (2012). Membrane tension maintains cell polarity by confining signals to the leading edge during neutrophil migration. Cell, 148:175–188.

[26] Howard, J., Grill, S., and Bois, J. (2011). Turing’s next steps: the mechanochemical basis of morphogenesis. Nat. Rev. Mol. Cell. Biol., 12:392–398.

[27] Inoue, T. and Meyer, T. (2008). Synthetic activation of endogenous PI3K and Rac identifies an AND-gate switch for cell polarization and migration. PLoS One, 3:e3068.

[28] Ip, Y. and Gridley, T. (2002). Cell movements during gastrulation: snail dependent and independent pathways. Curr. Opin. Genet. Dev., 12:423–429.

[29] Jilkine, A. and Edelstein-Keshet, L. (2011). A comparison of mathematical models for polarization of single eukaryotic cells in response to guided cues. PLoS Comput. Biol, 7:e1001121.

[30] Kan-On, Y. (1998). Bifurcation structure of stationary solutions of a Lotka-Volterra competition model with diffusion. SIAM J. Math. Anal., 29:424–436.

[31] Levchenko, A. and Iglesias, P. (2002). Models of eukaryotic gradient sensing: application to chemotaxis of amoebae and neutrophils. Biophys. J., 82:50–63.

[32] Lomakin, A., Lee, K., Han, S., Bui, D., Davidson, M., Mogilner, A., and Danuser, G. (2015). Competition for actin between two distinct F-actin networks defines a bistable switch for cell polarization. Nat. Cell Biol., 17:1435–1445.

[33] Ma, X., Daglyian, O., Hahn, K., and Danuser, G. (2018). Profiling cellular morphodynamics by spatiotemporal spectrum decomposition. PLoS Comput. Bio., 14:e1006321.

[34] Machesky, L. M. and Insall, R. H. (1998). SCAR1 and the related WiskottAldrich syndrome protein, wasp, regulate the actin cytoskeleton through the Arp2/3 complex. Curr. Biol., 8(25):1347 – 1356.

[35] Maiuri, P., Rupprecht, J., Wieser, S., Ruprecht, V., Benichou, O., Carpi, N., Coppey, M., Beco, S. D., Gov, N., Heisenberg, C., Crespo, C. L., Lautenschlaeger, F., Berre, M. L., Lennon-Dumenil, A.-M., Raab, M., Thiam, H.-R., Piel, M., Sixt, M., and Voituriez, R. (2015). Actin flows mediate a universal coupling between cell speed and cell persistence. Cell, 161:374–386.

[36] Maree, A., Jilkine, A., Dawes, A., Grieneisen, V., and Edelstein-Keshet, L. (2006). Polarization and movement of keratocytes: a multiscale modelling approach. Bull. Math. Biol., 68:1169–1211.

[37] McCaffrey, L. and Macara, I. (2012). Signaling pathways in cell polarity. Cold Spring Harb. Perspect. Biol., 4:a009654.

[38] Meinhardt, H. and Gierer, A. (1974). Applications of a theory of biological pattern formation based on lateral inhibition. J. Cell Sci., 15:321–346.

[39] Mogilner, A. and Keren, K. (2009). The shape of motile cells. Curr. Biol., 19:R762–771.

[40] Moissoglu, K. and Schwartz, M. (2014). Spatial and temporal control of Rho GTPase functions. Cell Logist., 4:e943618.

[41] Moissoglu, K., Slepchecko, B., Meller, N., Horwitz, A., and Schwartz, M. (2006). In vivo dynamics of Rac-membrane interactions. Mol. Biol. Cell, 17:2770–2779.

[42] Mori, Y., Jilkine, A., and Edelstein-Keshet, L. (2008). Wave-pinning and cell polarity from a bistable reaction-diffusion system. Biophys. J., 94:3684–3697.

[43] Mullins, R. (2010). Cytoskeletal mechanisms for breaking cellular symmetry. Cold Spring Harb. Perspect. Biol., 2:a003392.

[44] Munro, E., Nance, J., and Priess, J. R. (2004). Cortical flows powered by asymmetrical contraction transport PAR proteins to establish and maintain anterior-posterior polarity in the early *C. elegans* embryo. Dev. Cell, 7:413 – 424.

[45] Nguyen, T., Park, W., Park, B., Kim, C., Oh, Y., Kim, J., Choi, H., Kyung, T., Kim, C., Lee, G., Hahn, K., Meyer, T., and Heo, W. (2016). PLEKHG3 enhances polarized cell migration by activating actin filaments at the cell front. Proc. Natl. Acad. Sci. USA, 113:10091–10096.

[46] Pablo, M., Ramirez, S., and Elston, T. (2018). Particle-based simulations of polarity establishment reveal stochastic promotion of Turing pattern formation. PLoS Comput. Biol., 14:e1006016.

[47] Parent, C. and Devreotes, P. (1999). A cell’s sense of direction. Science, 284:765–770.

[48] Park, J., Holmes, W., Lee, S., Kim, H., Kim, D., Kwak, M., Wang, C., Edelstein-Keshet, L., and Levchenko, A. (2017). Mechanochemical feedback underlies coexistence of qualitatively distinct cell polarity patterns within diverse cell populations. Proc. Natl. Acad. Sci. USA, 114:e5750–e5759.

[49] Peglion, F. and Goehring, N. (2019). Switching states: dynamic remodelling of polarity complexes as a toolkit for cell polarization. Curr. Opin. Cell Biol., 60:121–130.

[50] Pertz, O., Hodgson, L., Klemke, R., and Hahn, K. (2006). Spatiotemporal dynamics of RhoA activity in migrating cells. Nature, 440:1069–72.

[51] Prager-Khoutorsky, M., Lichtenstein, A., Krishnan, R., Rajendran, K., Mayo, A., Kam, Z., Geiger, B., and Bershadsky, A. (2011). Fibroblast polarization is a matrix-rigidity-dependent process controlled by focal adhesion mechanosensing. Nat. Cell. Biol., 13:1457–1465.

[52] Ridley, A. (2006). Rho GTPases and actin dynamics in membrane protrusions and vesicle trafficking. Trends Cell Biol., 16:522–529.

[53] Ridley, A. J. (2001). Rho family proteins: coordinating cell responses. Trends Cell Biol., 11(12):471 – 477.

[54] Rotty, J. and Bear, J. (2014). Competition and collaboration between different actin assembly pathways allows for homeostatic control of the actin cytoskeleton. Bioarchitecture, 5:27–34.

[55] Schwartz, M. (2004). Rho signalling at a glance. J. Cell Sci., 117:5457–5458.

[56] Segal, D., Zaritsky, A., Schejter, E., and Shilo, B.-Z. (2018). Feedback inhibition of actin on Rho mediates content release from large secretory vesicles. J. Cell Biol., 217:1815.

[57] Srinivasan, S., Wang, F., Glavas, S., Ott, A., Hofmann, F., Aktories, K., Kalman, D., and Bourne, H. R. (2003). Rac and Cdc42 play distinct roles in regulating PIP3 and polarity during neutrophil chemotaxis. J. Cell Biol., 160(3):375–385.

[58] Su, W., Mruk, D., Wong, E., Lui, W., and Cheng, C. (2012). Polarity protein complex Scribble/Lgl/Dlg and epithelial cell barriers. Adv. Exp. Med. Biol., 763:149–170.

[59] Sun, Y., Do, H., Gao, J., Zhao, R., Zhao, M., and Mogilner, A. (2013). Keratocyte fragments and cells utilize competing pathways to move in opposite directions in an electric field. Curr. Biol., 23:569–574.

[60] Svitkina, T. (2018). The actin cytoskeleton and actin-based motility. Cold Spring Harb. Perspect. Biol., 10:a018267.

[61] Takeuchi, Y. (1989). Diffusion-mediated persistence in two-species competition Lotka-Volterra model. Math. Biosciences, 95:65–83.

[62] Tostevin, F. and Howard, M. (2008). Modeling the establishment of PAR protein polarity in the one-cell *C. elegans* embryo. Biophys. J., 95:45124522.

[63] Turing, A. (1952). The chemical basis of morphogenesis. Philos. Trans. R Soc. London Ser. B, 237:37–72.

[64] van der Gucht, J., Paluch, E., Plastino, J., and Sykes, C. (2005). Stress release drives symmetry breaking for actin-based movement. Proc. Natl. Acad. Sci. USA, 102:7847–7852.

[65] van Leeuwen, F., Kain, H., van der Kammen, R., Michiels, F., Kranenburg, O., and Collard, J. (1997). The guanine nucleotide exchange factor Tiam1 affects neuronal morphology; opposing roles for the small GTPases Rac and Rho. J. Cell Biol., 139:797–807.

[66] Vekhovsky, A., Svitkina, T., and Borisy, G. (1999). Self-polarization and directional motility of cytoplasm. Mol. Cell, 9:11–20.

[67] Weiner, O., Marganski, W., Wu, L., Altschuler, S., and Kirschner, M. (2007). An actin-based wave generator organizes cell motility. PLoS Biol., 9:e221.

[68] Weiner, O., Servant, G., Welch, W., Mitchison, T., Sedat, J., and Bourne, H. (1999). Spatial control of actin polymerization during neutrophil chemotaxis. Nat. Cell Biol., 1:75–81.

[69] Weiner, O. D. (2002). Rac activation: P-Rex1 – a convergence point for PIP3 and G*βγ*? Curr. Biol., 12(12):R429 – R431.

[70] Wong, K., Pertz, O., Hahn, K., and Bourne, H. (2006). Neutrophil polarization: spatiotemporal dynamics of RhoA activity support a self-organizing mechanism. Proc. Natl. Acad. Sci. USA, 103:3639–3644.

[71] Wu, C.-F., Chiou, J.-G., Minakova, M., Woods, B., Tsygankov, D., Zyla, T., and et al (2015). Role of competition between polarity sites in establishing a unique front. eLife, 4:e11611.

[72] Xu, J., Wang, F., van Keymeulen, A., Herzmark, P., Straight, A., Kelly, K., Takuwa, Y., Sugimoto, N., Mitchison, T., and Bourne, H. (2003). Divergent signals and cytoskeletal assemblies regulate self-organizing polarity in neutrophils. Cell, 114:201–214.

[73] Yam, P., Wilson, C., Ji, L., Hebert, B., Barnhart, E., Dye, N., Wiseman, P., Danuser, G., and Theriot, J. (2007). Actin-myosin network reorganization breaks symmetry at the cell rear to spontaneously initiate polarized cell motility. J. Cell Biol., 178:1207–1221.

[74] Yamada, S. and Nelson, W. (2007). Localized zones of Rho and Rac activities drive initiation and expansion of epithelial cell-cell adhesion. J. Cell Biol., 178:517–527.

[75] Zhang, B. and Zheng, Y. (1998). Negative regulation of Rho family GTPases Cdc42 and Rac2 by homodimer formation. J. Biol. Chem., 273:25728–25733.

## Supplemetary References

[1] Altschuler, S., Angenent, S., Wang, Y., and Wu, L. (2008). On the spontaneous emergence of cell polarity. Nature, 454:886–889.

[2] Nguyen, T., Park, W., Park, B., Kim, C., Oh, Y., Kim, J., Choi, H., Kyung, T., Kim, C., Lee, G., Hahn, K., Meyer, T., and Heo, W. (2016). PLEKHG3 *enhances polarized cell migration by activating actin filaments at the cell front*. Proc Natl Acad Sci U S A, 113:10091–10096.

[3] Neilson, M., Veltman, D., van Haastert, P., Webb, S., Mackenzie, J., and Insall, R. (2011). Chemotaxis: a feedback-based computational model robustly predicts multiple aspects of real cell behaviour. PLoS Biol., 9:e1000618.

[4] Weiner, O.D. (2002). Rac activation: P-Rex1 – a convergence point for PIP3 and Gβγ?. Curr Biol, 12(12):R429–431.

[5] Inoue, T. and Meyer, T. (2008). Synthetic activation of endogenous PI3K and Rac identifies an AND-gate switch for cell polarization and migration. PLoS One, 3:e3068.

[6] Byrne, K., Monsefi, N., Dawson, J., Degasperi, A., Bukowski-Wills, J., Volinsky, N., Dobrzynski, M., Birtwistle, M., Tsyganov, M., Kiyatkin, A., Kida, K., Finch, A., Carragher, N., Kolch, W., Nguyen, L., von Kriegsheim, A., and Kholodenko, B. (2016). Bistability in the Rac1, PAK, and RhoA signaling network drives actin cytoskeleton dynamics and cell motility switches. Cell Syst., 2:3848.

[7] Guilluy, C., Dubash, A., and Garcia-Mata, R. (2011). Analysis of RhoA and Rho GEF activity in whole cells and the cell nucleus. Nat. Protoc., 6:2050–2060.

[8] Burridge, K. and Wennerberg, K. (2004). Rho and Rac take center stage. Cell, 116:167179.

[9] Xu, J., Wang, F., van Keymeulen, A., Herzmark, P., Straight, A., Kelly, K., Takuwa, Y., Sugimoto, N., Mitchison, T., and Bourne, H. (2003). Divergent signals and cytoskeletal assemblies regulate self-organizing polarity in neutrophils. Cell, 114:201–214.

[10] van Leeuwen, F., Kain, H., van der Kammen, R., Michiels, F., Kranenburg, O., and Collard, J. (1997). The guanine nucleotide exchange factor Tiam1 affects neuronal morphology: opposing roles for the small GTPases Rac and Rho. J. Cell Biol., 139:797–807.

[11] Wang, Y., Ku, C., Zhang, E., Artyukhin, A., Weiner, O., Wu, L., and Altschuler, S. (2013). Identifying network motifs that buffer front-to-back signaling in polarized neutrophils. Cell Rep., 3:1607–1616.

[12] Alberts, B., Johnson, A., Lewis, J., Morgan, D., Raff, M., Roberts, K., and Walter, P., editors (2008). Intracellular membrane traffic. Garland Science, New York.

[13] Mogilner, A. and Keren, K. (2009). The shape of motile cells. Curr Biol., 19:R762–771.

[14] Mori, Y., Jilkine, A., and Edelstein-Keshet, L. (2008). Wave-pinning and cell polarity from a bistable reaction-diffusion system. Biophys J, 94:3684–3697.

[15] Das, S., Yin, T., Yang, Q., Zhang, J., Wu, Y., and Yu, J. (2015). Single-molecule tracking of small GTPase Rac1 uncovers spatial regulation of membrane translocation and mechanism for polarized signaling. Proc Natl Acad Sci U S A, 112:e267–e276.

[16] Zhang, B. and Zheng, Y. (1998). Negative regulation of Rho family GTPases Cdc42 and Rac2 by homodimer formation. J Biol Chem., 273:25728–25733.

[17] Moissoglu, K., Slepchecko, B., Meller, N., Horwitz, A., and Schwartz, M. (2006). In vivo dynamics of Rac-membrane interactions. Mol. Biol. Cell, 17:2770–2779.

[18] Falkenberg, C. and Loew, L. (2013). Computational analysis of Rho GTPase cycling. PLoS Comput Biol., 9:e1002831.

